# Adaptive radiation and burst speciation of hillstream cyprinid fish *Garra* in African river

**DOI:** 10.1101/2021.05.04.442598

**Authors:** Boris Levin, Evgeniy Simonov, Paolo Franchini, Nikolai Mugue, Alexander Golubtsov, Axel Meyer

## Abstract

Adaptive radiation of fishes was long thought to be possible only in lacustrine environments. Recently, several studies have shown that also riverine and stream environments provide the ecological opportunity for adaptive radiation. In this study, we report on a riverine adaptive radiation of six ecomorphs of cyprinid hillstream fishes of the genus *Garra* in a river located in the Ethiopian Highlands in East Africa. *Garra* are predominantly highly specialized algae-scrapers with a wide distribution ranging from Southeastern Asia to Western Africa. However, adaptive phenotypic diversification in mouth type, sucking disc morphology, gut length and body shape have been found among these new species in a single Ethiopian river. Moreover, we found two novel phenotypes of *Garra* (‘thick-lipped’ and ‘predatory’) that were not described before in this species-rich genus (>160 species). Mitochondrial and genome-wide data suggest monophyletic, intra-basin evolution of *Garra* phenotypic diversity with signatures of gene flow from other local populations. Although sympatric ecomorphs are genetically distinct and can be considered to being young species as suggested by genome-wide SNP data, mtDNA was unable to identify any genetic structure suggesting a recent and rapid speciation event. Furthermore, we found evidence for a hybrid origin of the novel ‘thick-lipped’ phenotype, as being the result of the hybridization of two other sympatrically occurring species. Here we highlight how, driven by ecological opportunity, an ancestral trophically highly specialized lineage is likely to have rapidly adaptively radiated in a riverine environment, and that this radiation was promoted by the evolution of novel feeding strategies.

## Introduction

Unravelling the mechanisms underpinning the biological diversity remains a major challenge in evolutionary biology. With more than 28,000 species, teleost fishes are the most diverse lineage of vertebrates, and thus an ideal system to address questions regarding diversification. The stunning phenotypic diversity of bony fishes has largely been produced through the process of adaptive radiation, the rapid proliferation of multiple ecologically distinct species from a common ancestor (Schluter, 2000). One of the most extraordinary examples of both adaptive radiation and explosive diversification is represented by the cichlid fishes inhabiting the East African Great Lakes (Kocher, 2004). According to Losos (2010) and Givnish (2015) adaptive radiation and explosive diversification are distinct phenomena: the former may or may not result in, or be accompanied by the latter. The evolutionary success of the cichlids, unmatched among vertebrates, has been promoted by a combination of different factors, where a dominant role has been played, for example, by limited dispersal (because of territoriality and mouth-brooding) and sexual selection for nuptial coloration and mating behavior (Henning & Meyer, 2014; Meyer, Kocher, Basasibwaki, & Wilson, 1990; Seehausen, 2000; Wagner, Harmon, & Seehausen, 2012). It has been suggested, however, that trophic radiation had preceded the diversification driven by other factors at least in cichlids of Lake Tanganyika (Muschick et al., 2014), a cradle of all other East African haplochromine radiations (Salzburger, Mack, Verheyen, E., & Meyer, 2005). Adaptive radiations and diversification bursts were found not only in cichlids, but also in other fish groups, even though in smaller scale, and often in a parallel manner - coregonids, Arctic charrs, and sticklebacks (e.g. Broderson, Post, & Seehausen, 2018; DeFaveri & Merila, 2013; Jacobs et al., 2020; McKinnon & Rundle, 2002; Præbel et al., 2013; Peichel et al., 2001; Schluter, 2000; Skúlason, 1999; Terekhanova et al., 2014) - some of the best known examples of intralacustrine radiations.

The most supported cases of monophyletic, closely related fish species that are believed to have arisen through an adaptive radiation event have been described from lakes rather than rivers (Meyer et al. 1990; Seehausen, 2006; Sturmbauer, 1998; Taylor, 1999). For long time, riverine environment has not been considered suitable for adaptive radiation because of its unstable hydrological regimes, reduced habitat diversity and the commonly shallow and narrow watercourses that might facilitate gene flow (Seehausen & Wagner, 2014). However, during the last two decades, examples of fish adaptive radiations occurring in rivers have been reported (Burress et al., 2018; Dimmick, Berendzen, & Golubtsov, 2001; Levin, Simonov, Dgebuadze, Levina, & Golubtsov, 2020; Melnik, Markevich, Taylor, Loktyushkin, & Esin, 2020; Piálek, Říčan, Casciotta, Almirón, & Zrzavý, 2012; Schwarzer, Misof, Ifuta, & Schliewen, 2011; Whiteley, 2007). Although several cases of riverine diversification of cichlid fishes are considered as remnants of adaptive radiations occurred in the palaeo-Lake Makgadikgadi before it dried up back in the Holocene (Joyce et al., 2005), mounting evidence suggests that some fish species flocks of other species than cichlids have diversified within rivers (Burress et al., 2018; Levin et al., 2019; 2020; Melnik et al., 2020; Piálek et al., 2012)

In the present study, we investigate a highly diverse fish group that presumably adaptively radiated in riverine environments. The genus *Garra* is a species-rich lineage of labeonine cyprinids comprising more than 160 species and is distributed from Southeast Asia to West Africa (Fricke, Eschmeyer, & Van der Laan, 2021; Yang et al., 2012). *Garra* are mostly moderate-sized fish (usually less than 20 cm in length) with sucking gular disc that inhabit the rhithron zone of river systems (Kottelat, 2020). They are predominantly highly specialized algae scrapers that graze periphyton from rocks and stones using widened jaws equipped with horny scrapers. However, adaptations to still waters such as caves or lacustrine environment have been documented in the *Garra*, although rarely, accompanied by a reduction of the gular disc and a change of the foraging strategy from algae scraping to planktivory (Geremew, 2007; Kottelat, 2020; Segherloo et al., 2018; Stiassny & Getahun, 2007; www.briancoad.com).

The Ethiopian Highlands are recognized as a center of *Garra* diversity within Africa (Golubtsov, Dgebuadze, & Mina, 2002; Stiassny & Getahun, 2007), where 13 described species out of the total 23 found in Africa are recorded (Moritz, El Dayem, Abdallah, & Neumann, 2019). An assemblage of six *Garra* ecomorphs exhibiting extreme morphological diversity was recently discovered in the Sore River (the White Nile Basin) in southwestern Ethiopia during a survey of the Ethiopian fishes (Golubtsov, Cherenkov, & Tefera, 2012). In particular, two of the six forms display features not found elsewhere within the generic range: a form with a pronounced predatory morphology (large-sized, large-mouthed, with reduced sucking disk and a short gut that is equal to body length) and one with ‘rubber’ lips and prolonged snout region (Fig. 1, Table 1). The other four forms from the Ethiopian *Garra* assemblage drastically differ in mouth and gular disc morphology as well as in body shape (Fig. 1).

**Fig. 1A.**
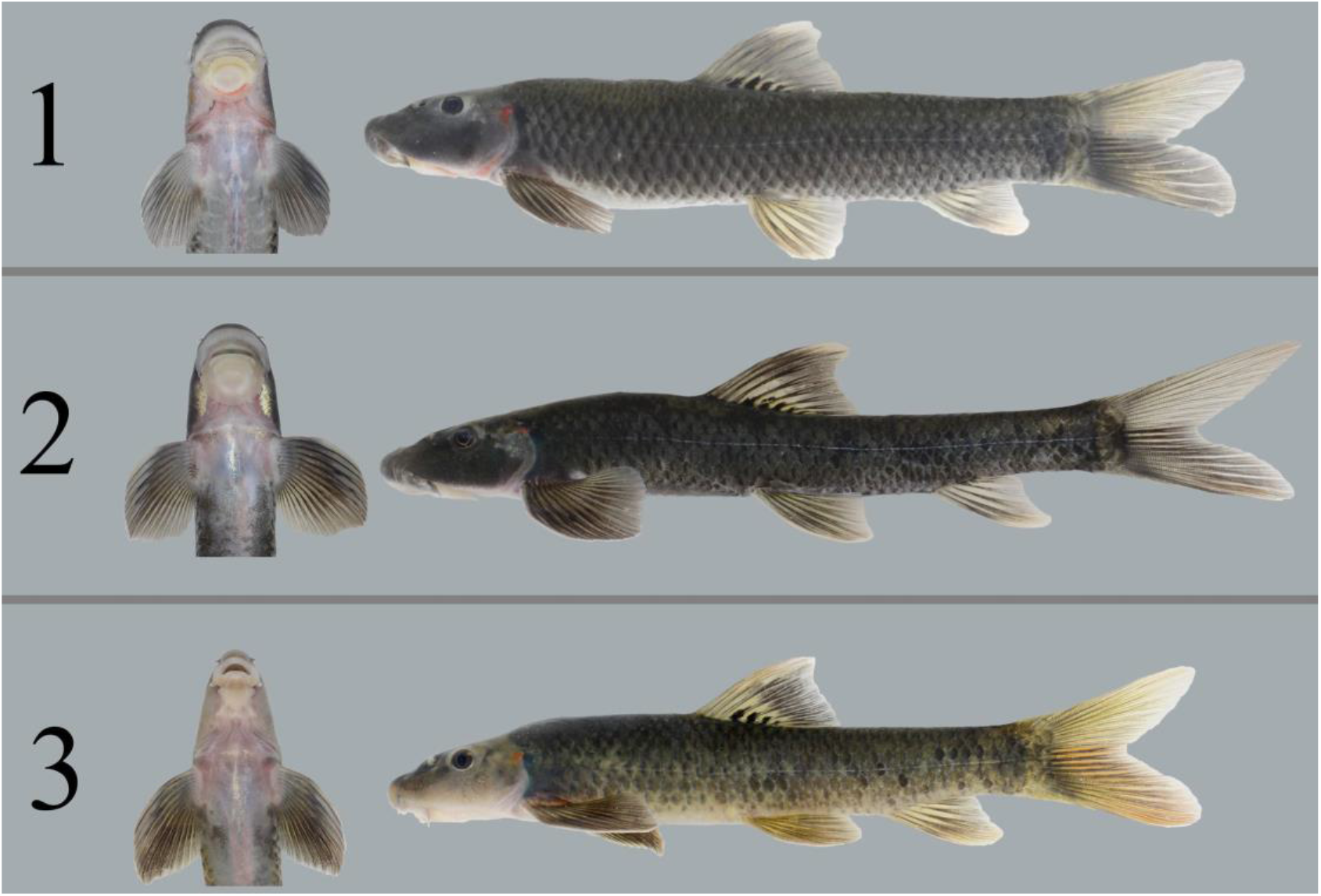
*Garra* ecomorphs 1-3 from the Sore River: 1 - ‘generalized’: 136 mm SL; 2 - ‘stream-lined’: 99 mm SL; 3 - ‘narrow-mouth’: 100 mm SL.

**Fig. 1B.**
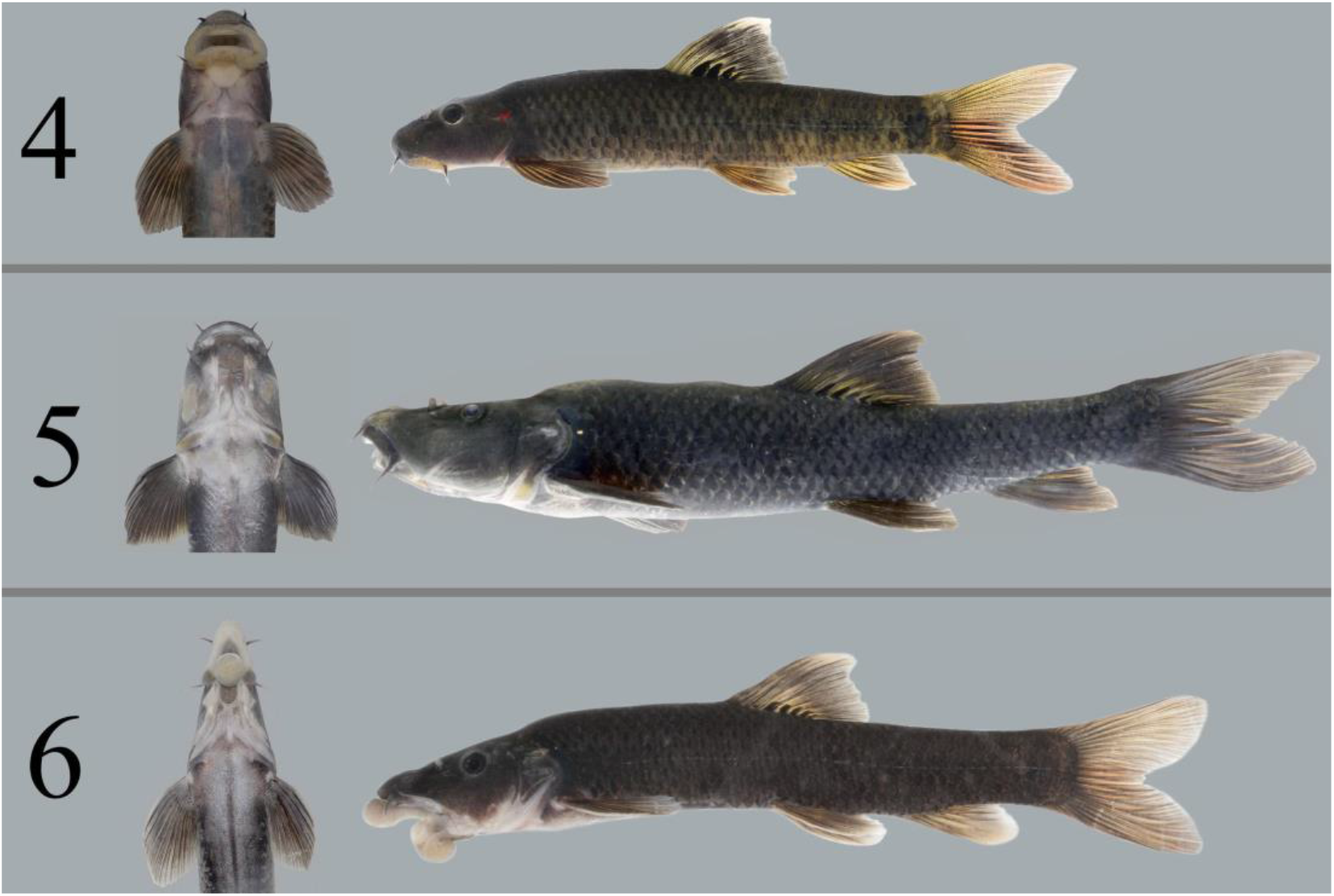
*Garra* ecomorphs 4-6 from the Sore River: 4 - ‘wide-mouth’: 100 mm SL; 5 - ‘predator’: 193 mm SL; 6 - ‘thick-lipped’: 128 mm SL.

**Table 1.** Common names of the six ecomorphs of African *Garra* from the Sore River, and the preliminary qualitative descriptions used in the field to identify each form.

| Name used in the text | Basal description |
| --- | --- |
| No. 1, ‘generalized’ | Well-developed round-shaped gular disc of type C with free posterior margin (disc classification follows Stiassny & Getahun, 2007). Body shape is generalized for *Garra*. |
| No. 2, ‘stream-lined’ | Slender stream-line body with slim caudal peduncle and increased pectoral fins. Disc of type C. |
| No. 3, ‘narrow-mouth’ | Disc is reduced in size, elongated, oval-shaped (closer to type A). Narrow mouth often with groove on lower jaw. |
| No. 4, ‘wide-mouth’ | Disc is reduced in size, triangle-shaped. Wide mouth with significantly enlarged labellum (sensu Kottelat, 2020). Disc of type B in degree of development. |
| No. 5, ‘predator’ | Completely or almost completely reduced gular disc (type A when presented). Wide head and mouth. This ecomorph achieves larger size compared to others. Largest individuals have nuchal hunch and almost terminal mouth with a bony projection on the lower jaw and matching incision on the upper jaw. |
| No. 6, ‘thick-lipped’ | Greatly developed lips, referred to as ‘rubber lips’ (Matthes, 1963). Intermediate lobe of the lower lip is ball-shaped and unattached. Gular disc is greatly reduced, oval-shaped (type A). Only two individuals recorded. |

Our goals were twofold: i) to investigate the morpho-ecological relationships of six *Garra* sympatric ecomorphs from the Sore River, and ii) to test whether this assemblage has evolved sympatrically. In detail, we aimed at elucidating the population structure and evolutionary history of these ecomorphs using both mitochondrial DNA (mtDNA, cytochrome *b*) and genome-wide nuclear loci obtained with a double digest restriction-site associated DNA (ddRAD) approach.

## Materials and Methods

### Study area

The Sore River is a headwater tributary of the Baro-Akobo-Sobat drainage in the White Nile basin, (south-western Ethiopia, northern East Africa). It drains the Ethiopian Highlands close to the south-western escarpment. The region is covered by moist Afromontane forest that is drastically shrinking in the last decades due to agricultural development (Dibaba, Soromessa, & Workineh, 2019). The Sore is a rather little river with a length of *ca*. 160 km, its catchment area is *ca*. 2000 km^2^ and characterized by substantial seasonal variation of rainfall (dry season from December to March) (Kebede, Diekkrüger, & Moges, 2014). In comparison, the Italian Tiber River length is 406 km, its catchment area is 17375 km^2^ (https://en.wikipedia.org/wiki/Tiber). Elevation difference between the Sore source (altitude of ca. 2215 m asl, above sea level) and its confluence with the Gabba (Geba) River (alt. 963 m asl) is 1.25 km. The Sore River basin shares drainage boundaries with two of six major watersheds of Ethiopia: Blue Nile in the north-east and Omo-Turkana in the south-east.

We sampled the middle reaches of the Sore River at two sites: (1) at the City of Metu (8°18’42’’ N 35°35’54″ E, alt. 1550 m asl) and (2) ca. 35 km downstream along the river course (8°23’56″ N 35°26’18″ E, alt. 1310 m asl). The river width at the rapids sampled was 20-40 m at the beginning of the rainy season, depth <1 m, bottom consisted of rocks and large boulders. Fish fauna of the river segment under consideration includes (apart from *Garra* spp.) a species flock of *Labeobarbus* (Levin et al., 2020), *Enteromius* cf. *pleurogramma* (Boulenger 1902), *Labeo* cf. *cylindricus* Peters 1852, *Labeo forskalii* Rüppell 1835, *Chiloglanis* cf. *niloticus* Boulenger 1900 (at the lower site only), and introduced *Coptodon zillii* (Gervais 1848). Presence of the stony loach (*Afronemacheilus*) reported by Getahun and Stiassny (1998) from the Sore River at Metu could no longer be confirmed (Melaku, Abebe Getahun, & Wakjira, 2017; Prokofiev & Golubtsov, 2013; present study). Attempts to re-sample a stony loach by intensive electrofishing in 2012 have resulted in the discovery of the enormous morphological *Garra* diversity in the Sore River (Golubtsov et al., 2012). A hundred kilometers westward, from the lowland part (alt. ca. 500 m asl) of the same river drainage >100 fish species are recorded (Golubtsov & Darkov, 2008; Golubtsov, Darkov, Dgebuadze, 1995;) and >115 species from the Sudd and White Nile in Sudan and South Sudan (Moritz et al., 2019; Neumann, Obermaier, & Moritz, 2016;).

### Sampling

*Garra* samples from the Sore River were collected using a battery driven electrofishing device (LR-24 Combo Backpack, Smith-Root, USA), cast and frame nets in June 2012 and April 2014. In 2011-2014 comparative *Garra* samples were collected from nine sites in six main Ethiopian basins (Fig. 2, Table S1). Fish sampling was conducted under the umbrella of the Joint Ethiopian-Russian Biological Expedition (JERBE) with the permissions of National Fisheries and Aquatic Life Research Center (NFALRC) under Ethiopian Institute of Agricultural Research (EIAR) and Ethiopian Ministry of Science and Technology (presently Ministry of Innovation and Technology). Fish were killed with an overdose of an anesthetic MS-222, first preserved in 10% formalin and then transferred to 70% ethanol. From each specimen fin tissue samples were fixed with 96% ethanol. Some fish specimens were pictured using a Canon EOS 50D camera. All specimens (Supplementary Table S1) are deposited at the A.N. Severtsov Institute of Ecology and Evolution, at the Russian Academy of Sciences, Moscow, under provisional labels of JERBE.

**Fig. 2.**
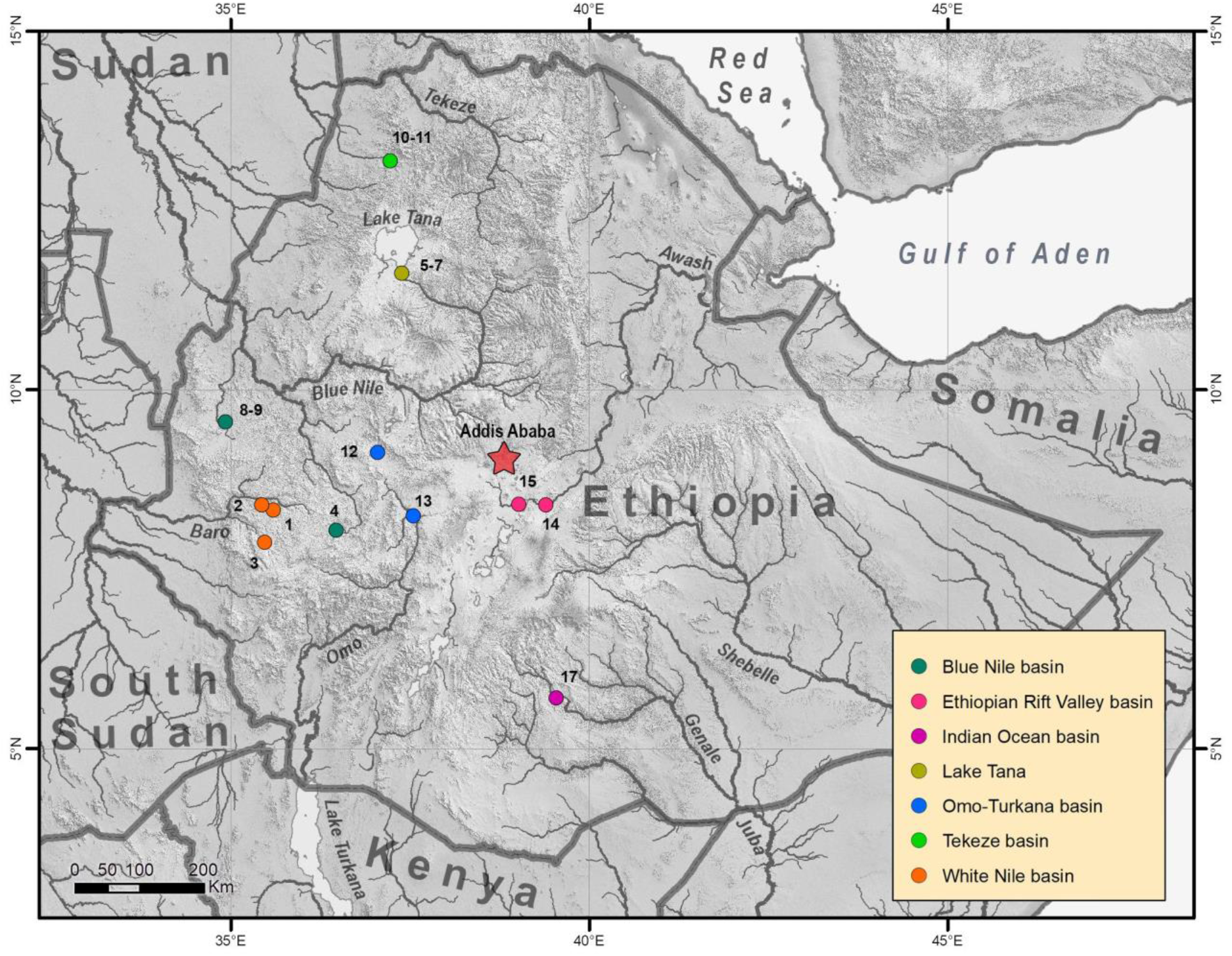
Sampling sites of *Garra* in Ethiopian Highlands and Ethiopian Rift Valley; loc. 1-2 are in the Sore River.

### Morphological analysis

#### Morphometry

The 28 morphometric characters from 107 individuals of all ecomorphs from the Sore River were examined following Hubbs and Lagler (1958) with additions from Menon (1964): standard length (SL), head length (HL), snout length (R), eye diameter (O), postorbital distance (PO), interorbital distance (IO), head width (HW), head height at nape (HH), head height at mid-of-eye (Hh), mouth width (MW), disc length (DL), disc width (DW), maximal body height (H), minimal body height at caudal peduncle (h), predorsal length (PL), postdorsal length (PDL), prepelvic length (PPL), preanal length (PAL), caudal peduncle length (CPD), dorsal fin base length (DFL), dorsal fin depth (DFP), anal fin base length (AFL), anal fin depth (AFD), pectoral fin length (PFL), ventral fin length (VFL), pectoral-ventral fin distance (PV), ventral-anal fin distance (VA), and distance between anal opening and anal fin (DAA). Measurements were done using a digital caliper (to nearest 0.1 mm). All measurements were performed by one operator for the purpose of consistency as recommended by Mina, Levin, and Mironovsky (2005).

Measured individuals had body length varied from 43.6 to 185.0 mm SL: ecomorph 1 (71.5-151.0), ecomorph 2 (70.9-160.2), ecomorph 3 (49.3-100.6), ecomorph 4 (49.3-90.6), ecomorph 5 (43.6-81.0; one individual had outstanding length - 185.0), ecomorph 6 (118.4; 139.4) (defined as in Fig. 1 and Table 1), intermediate phenotypes (59.3-105.2). The proportions of head and body were used for principal component analysis (PCA) - measurements of head parts were divided for head length and measurements of body parts were divided for standard length. Data was scaled. The gular disc in some specimens of ecomorph 5 was greatly reduced which hampered the detection of its borders. For the purpose of justification of the values of this character, the identical intermediate values were arbitrarily assigned for all specimens of this ecomorph. PCA was done using *prcomp* script implemented in R with a variance-covariance matrix.

#### Gut length and preliminary assay of a diet

Intestines were taken out from the body cavity of 62 preserved specimens of all ecomorphs except for no. 6 (represented by only two specimens), and measured using a ruler to the nearest 1 mm. The sample size for each ecomorph is provided in Table 2. The standard length (SL) of examined individuals varied from 40 to 131 mm, one individual of ecomorph 5 had outstanding length - 185 mm. The ratio of gut length (GL) to SL was used for subsequent analyses. The Kruskall-Wallis test for multiple independent samples with Benjamini-Hochberg method of control of false discovery rate (FDR) (Benjamini & Hochberg, 1995) of *p*-value was applied to check a significance of differences at p<0.05. The dependence of GL on SL was visualized using scatterplots and regressions. R-packages *ggplot2* and *PMCMR* were used to create plots and to test statistical significance of differences.

**Table 2.**
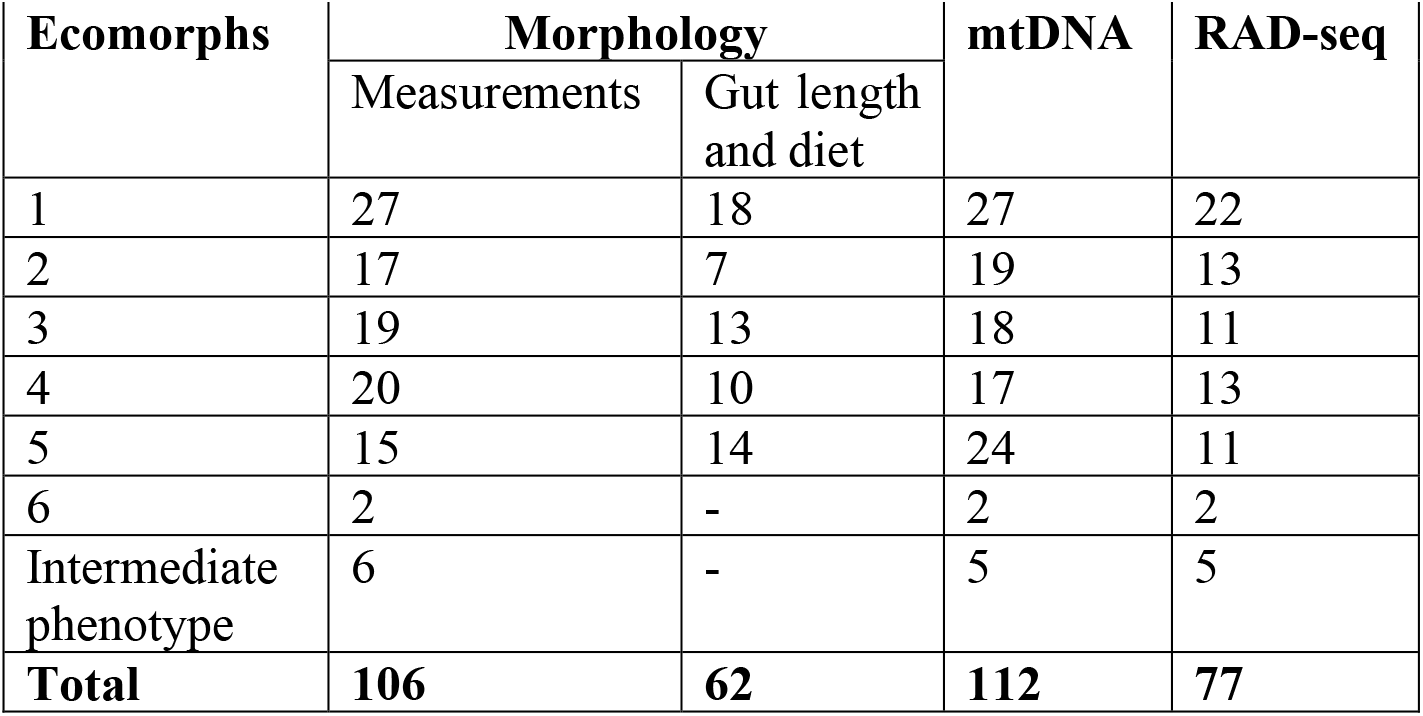
DNA and morphology sample numbers of *Garra* ecomorphs from the Sore River.

| Ecomorphs | Morphology |  | mtDNA | RAD-seq |
| --- | --- | --- | --- | --- |
|  | Measurements | Gut length and diet |  |  |
| 1 | 27 | 18 | 27 | 22 |
| 2 | 17 | 7 | 19 | 13 |
| 3 | 19 | 13 | 18 | 11 |
| 4 | 20 | 10 | 17 | 13 |
| 5 | 15 | 14 | 24 | 11 |
| 6 | 2 | - | 2 | 2 |
| Intermediate phenotype | 6 | - | 5 | 5 |
| **Total** | **106** | **62** | **112** | **77** |

Diet was assessed for the same individuals, whose intestine length was measured. The main ecological and systematic groups were registered using stereo-microscope Micromed MC-2-ZOOM and microscope Olympus CX41. A composite measure of diet, an index of relative importance, IRI (Hart, Calver, & Dickman, 2002), was used to assess contribution of different components to a diet. The diet components were grouped in several items i) periphyton, ii) benthos, iii) macrophytes, and iv) others.

#### DNA sampling, extraction, amplification, and sequencing - mtDNA data

DNA samples (n=107) were collected from *Garra* inhabiting the Sore River near the City of Metu in 2012 and 2014 from all six forms (see Table 2 for details). For comparison additional DNA samples (n=20) were collected from 8 *Garra* species inhabiting all main drainages of Ethiopia (10 localities – see map of sampling in Fig. 2). Total genomic DNA was extracted from ethanol-preserved fin tissues using the BioSprint 15 kit for tissue and blood (Qiagen). Sequences of the mitochondrial gene, cytochrome *b* (*cytb*) of 989 bp length, were amplified (see PCR conditions in Supplementary Material S2; Palumbi, 1996; Perdices & Doadrio, 2001). PCR products were visualized on 1% agarose gels, purified with ExoSAP-IT™ and sequenced at the Papanin Institute of Biology of Inland Waters (Russian Academy of Sciences) using an ABI 3500 sequencer. All new sequences were deposited in GenBank (Accession Numbers: xxx -will be provided upon acceptance, see Supplementary Table S1).

#### Analysis of mtDNA data

All sequences were aligned and edited using the MUSCLE algorithm (Edgar, 2004) as implemented in MEGA 6.0 (Tamura, Stecher, Peterson, Filipski, & Kumar, 2013). A final set that includes also comparative material from Genbank (African and non-African *Garra* as well as outgroups) encompassed 143 *cytb* sequences (https://www.ncbi.nlm.nih.gov) (Table S1). *Akrokolioplax bicornis* and *Crossocheilus burmanicus* were included as outgroups according to previously published phylogenies (Yang et al., 2012).

Gene tree reconstruction was performed using both maximum-likelihood (ML) and Bayesian inference (BI) approaches. Prior to these analyses all sequences were collapsed into common haplotypes using ALTER software (Glez-Peña, Gómez-Blanco, Reboiro-Jato, Fdez-Riverola, & Posada, 2010). We determined the best fit models of nucleotide substitution for each codon position of *cytb* and optimal partitioning scheme using either ModelFinder (as implemented in IQ-TREE 1.6.12; Kalyaanamoorthy, Minh, Wong, Von Haeseler, & Jermiin, 2017; Nguyen, Schmidt, Von Haeseler, & Minh, 2015) or PartitionFinder 2.1.1 (Lanfear, Calcott, Ho, & Guindon, 2012) under Bayesian Information Criterion (BIC). The partition scheme selected by ModelFinder (codon position 1 - K2P+R2; codon position 2 - HKY+F+I; codon position 3 - TN+F+G4) was subsequently used in ML search with IQ-TREE, using 1 000 bootstrap replicates.

Bayesian phylogenetic inference (BI) was carried out in MrBayes v. 3.2.6 (Ronquist et al., 2012). The selected partition scheme was following: codon position 1 with K80+I+G, codon position 2 with HKY+I, and codon position 3 with GTR+G. Two simultaneous analyses were run for 10^7^ generations, each with four MCMC chains sampled every 500 generations. Convergence of runs was assessed by examination of the average standard deviation of split frequencies and the potential scale reduction factor. In addition, stationarity was confirmed by examining posterior probability, log likelihood, and all model parameters by the effective sample sizes (ESSs) in the program Tracer v1.6 (Rambaut, Suchard, Xie, & Drummond, 2014). The gene trees resulting in ML and BI analyses were visualized and edited using FigTree v.1.4.4 (Rambaut, 2014). A haplotype network was constructed using the median joining algorithm (Bandelt, Forster, & Röhl, 1999) in PopArt 1.7 (Leigh & Bryant, 2015).

#### ddRAD-seq library preparation

High molecular weight DNA was isolated from fin tissue preserved in ethanol using QIAamp DNA Mini Kit (Qiagen, Germany) or obtained by purification of salt method extracted DNA (Aljanabi & Martinez, 1997) using CleanUp Standard kit (Evrogen, Moscow). The dsDNA quantity was measured using dsDNA HS Assay Kit for fluorometer Qubit 3 (Life Technologies, USA). ddRAD-library was constructed following the quaddRAD protocol (Franchini, Monné Parera, Kautt, & Meyer, 2017) using restriction enzymes *Pst*I and *Msp*I. In total, 77 DNA samples of *Garra* ecomorphs from the Sore River (see Table 2) and 11 DNA samples from five other species of Ethiopian *Garra* from adjacent basins were sequenced by two independent runs of Illumina HiSeq2500 and Illumina X Ten (2 x 150 bp paired-end reads). The raw sequencing data were demultiplexed by the sequencing provider using outer Illumina TruSeq dual indexes.

#### Processing of RAD-seq data

The resulting reads were trimmed for remaining adapters and low quality reads Cutadapt implemented in the Trim Galore 0.4.5 package (https://github.com/FelixKrueger/TrimGalore - Martin, 2011). Read quality was assessed with FastQC 0.11.7 (Andrews & Krueger, 2010) and MultiQC 1.7 (Ewels, Magnusson, Lundin, & Käller, 2016) before and after trimming. Further demultiplexing of individually barcoded samples, construction and cataloging of RAD-loci, and SNP calling were done with STACKS 2.41 package (Catchen, Hohenlohe, Bassham, Amores, & Cresko, 2013). Identification and removal of PCR duplicates were done using the *‘clone_filter’* module of STACKS). STACKS module *‘process_radtags’* was used to demultiplex reads by the dual index inner barcodes and obtain separate fastq files for each individual. Samples that failed to produce more than 100 000 reads were excluded from further processing. To additionally evaluate data quality and identify possible contaminated samples, the reads were mapped to the reference genome of common carp *Cyprinus carpio* (GCF_000951615.1) using bowtie2 2.3.5 (Langmead & Salzberg, 2012) with ‘--local-sensitive’ presettings. Then, only Read 1 (R1) files were used for downstream processing and analyses. Prior to next steps, these R1 reads were trimmed at their 3̀ ends to a uniform length of 130 bp to reduce the influence of sequencing error (due to declined base quality at 3̀ end).

The *de novo* pipeline of STACKS was used to assemble loci and perform genotype calling. We selected optimal parameters using the approach suggested by Paris, Stevens, & Catchen (2017). Following the aforementioned procedure, we found that minimum stack depth (*-m*) of 5, distance allowed between stacks (-*M*) of 3, and the maximum distance required to merge catalog loci (-*n*) of 5 provided the best balance between data quality and quantity for our dataset (Fig. S1).

#### Population genomic analyses

Individual genotypes of sympatric *Garra* ecomorphs from the Sore River were exported to a vcf file using the ‘*populations*’ module of STACKS with the following settings: (i) loci genotyped in at least 90% of samples (-r 0.90) were kept; (ii) SNPs with a minor allele frequency (--min-maf) less than 0.04 and a maximum observed heterozygosity (--max_obs_het) above 0.99 were pruned; (iii) only single SNP per RAD locus was retained, to avoid inclusion of closely linked SNPs. We applied VCFtools 0.1.16 (Danecek et al., 2011) for further filtering of the dataset based on mean coverage and fraction of missing data for each sample. Samples with more than 20% of missing data were blacklisted and excluded from further analyses. Thus, a high-quality dataset of 679 SNPs and 77 individuals was obtained and used for downstream population genetics analyses.

First, Principal Component Analysis (PCA) was performed using the ‘*glPca’* function of the R-package *adegenet* 2.1.1 (Jombart, 2008; Jombart & Ahmed, 2011). Next, *rmaverick* 1.0.5 (former MavericK; Verity & Nichols, 2016) was used to infer population structure. This program estimates evidence for different numbers of populations (*K*), and different evolutionary models via generalised thermodynamic integration (GTI). A range of *K* values between 1 and 10 were explored, using 300 000 burn-in MCMC iterations and 10 000 sampling iterations. Convergence of MCMC was automatically tested every 1 000 burn-in iterations by activating option ‘auto_converge’. This allows exit burn-in iterations when convergence is reached and immediately proceeds to sampling iterations. Parameter ‘rungs’ was set to 10 (number of multiple MCMC chains with different ‘temperature’ to run simultaneously). Both no admixture and admixture models were run, and compared by plotting values of the posterior distribution and overall model evidence in log space (log-evidence) (Fig. S2-S5). According to this comparison, the admixture model is decisively supported over the no admixture model, and used here to report the results. The same protocol was followed for consecutive hierarchical *rmaverick* runs for the identified clusters. Finally, global and pairwise Reich-Patterson F_ST_ values (Reich, Thangaraj, Patterson, Price, & Singh, 2009) with respective 95% confidence intervals for ecomorphs/genetic clusters were calculated using the R script from Junker et al. (2020). Basic genetic diversity statistics were calculated using the ‘*populations*’ module of STACKS.

To test for the gene flow between ecomorphs\genetic clusters, we used the Patterson’s D statistic (ABBA-BABA test), along with the ƒ_4_-ratio statistic (Patterson et al., 2012) and its ƒ-branch metric (Malinsky et al., 2018), as implemented in Dsuite 0.4 software package (Malinsky, Matschiner, & Svardal, 2021). Patterson’s D statistic is a widely used and robust tool to detect introgression between populations or closely related species, and to distinguish it from incomplete lineage sorting (ILS). The ƒ_4_-ratio statistic is a similar method aiming to estimate an admixture fraction. The ƒ-branch metric is based on ƒ_4_-ratio results and serves to assign gene flow evidence to specific branches on a phylogeny. These tests were performed on a group containing ecomorphs\genetic clusters 2b, 3, 4, and 6, while the rest were used as outgroup (in accordance with the results of our phylogenomic analysis).

#### Phylogenomic analyses

IQ-TREE 2.0.5 (Minh et al., 2020) was used for ML phylogenetic analyses of RAD-seq data. First dataset included one to three specimens of each *Garra* ecomorph from the Sore river and other Ethiopian *Garra* species from adjacent basins. Multiple sequence alignments of all loci and respective partition files were created using the ‘--phylip-var-all’ option of ‘*populations*’ module of STACKS package. Heterozygous sites within each individual were encoded using IUPAC notation. During the analysis each RAD-locus was treated as a separate partition with independent best-fit substitution model. Node support values were obtained using ultrafast bootstrap procedure (Hoang, Chernomor, von Haeseler, Minh, & Vinh, 2018) with 1 000 replicates. We also used SVDQuartets algorithm (Chifman & Kubatko, 2014) as implemented in PAUP* 4.0a168 (Swofford, 2003) to perform species-tree inference under the multi-species coalescent model using 18,988 SNPs (single random SNP per locus, minor allele frequency cutoff 0.04, maximum observed heterozygosity cutoff: 0.99). Node support was estimated with 1 000 bootstrap replicates.

The second dataset consisted of all genotyped specimens of sympatric *Garra* ecomorphs from the Sore River and a single, most closely related outgroup (*G.* cf. *dembeensis* from the Barokalu River, as revealed by the analysis of the first phylogenomic dataset that included samples from all the localities in Figure 2). It was analysed with IQ-TREE as described above, except for GTR+G substitution model was used for each partition. The phylogenetic trees were visualized and edited using FigTree 1.4.4 (Rambaut & Drummond, 2008).

## Results

### Trophic Morphology

PCA of head and body proportions of six sympatric ecomorphs from the Sore River revealed five well-defined clusters (Fig. 3A). Four clusters represent ecomorphs 3, 4, 5, and 6, while the fifth includes individuals from ecomorphs 1 and 2. The ecomorph 5 is the most divergent. PC1 explained 72.3% of the total variance, while PC2 10.2%. The eigenvector with the highest eigenvalues for PC1 were head proportions - nine of ten most loaded ones (especially gular disc proportions, mouth width, interorbital distance, and snout length). The same pattern was detected for PC2 - nine of ten most loaded characters belonged to head proportions (mainly disc length, mouth width, height of head at nape and at eyes etc. - see Table S2 for details).

**Fig. 3.**
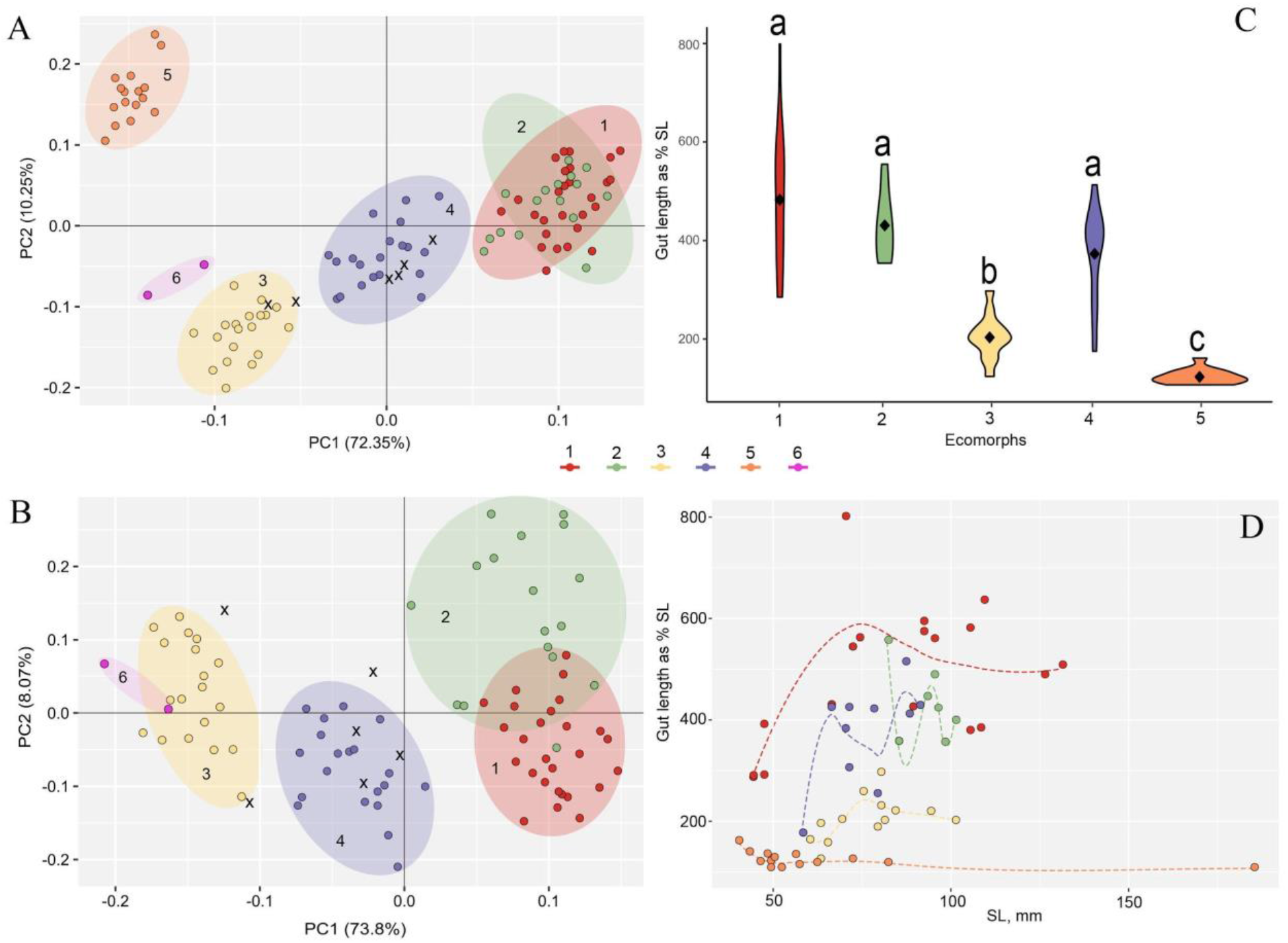
(A) PCA of body and head proportions of six sympatric ecomorphs from the Sore River (n=107); (B) PCA of body and head proportions of five sympatric ecomorphs from the Sore River (n=90) excluding the most divergent sample, ecomorph 5. X designates intermediate phenotypes; (C) Gut length of five sympatric *Garra* ecomorphs from the Sore River represented as violin boxplots. Middle points are the means, and the box show the range respectively, samples are combined and each contains between 7 (ecomorph 2) and 18 (ecomorph 1) individuals, for a total of 62 individuals. Different lowercase letters above the boxplots indicate significant differences between ecomorphs (*p* < 0.05, Kruskal-Wallis test with BH adjustment of *p*-value); (D) Dependence of gut length on body length in five *Garra* ecomorphs from the Sore River with smooth local regression lines (Loess regression).

After excluding ecomorph 5, the ecomorphs 1 and 2 became more distinguishable with low overlapping (Fig. 3B). The PC1 explained 73.8% of variance, while PC2 8.1%. The most loaded eigenvectors of both PC1 and PC2 were from head proportions with few more contributions of some body proportion characters (see Table S3). The difference between ecomorphs 1 and 2 revealed in PC2 is explained by height of head at both nape and eyes, interorbital distance, head width, body height as well as other characters (Table S3).

### Gut length and preliminary data on diet

Gut length broadly varied consistently between ecomorphs (Fig. 3C). Shortest guts (107-160 % SL) were detected in ecomorph 5 suggested a predatory trophic type, while the longest guts were recorded in ecomorphs 1 (285-799 % SL) and 2 (354-555 % SL) that possessed the well-developed gular disc and therefore are specialized algal grazers, as also shown by their gut contents (see below). Other ecomorphs had intermediate values gut lengths: ecomorph 3 - 124-295 % SL, and ecomorph 4 - 175-513 % SL, respectively. Broad intra-group variation is explained by increase of gut length with body length detected in some ecomorphs (Fig. 3D). Nevertheless, the similar-sized individuals are divergent in gut length at the same manner that presented in Fig. 3C. Ecomorph 5 having the shortest gut displays even a slight decrease of gut length ontogenetically that was previously reported for piscivorous mode of feeding among African cyprinids (Levin et al., 2019).

The preliminary inspection of gut content revealed differences in the diet between some ecomorphs. Ecomorphs 1 and 2 had permanently filled intestines full of periphyton (diatom, green, and charophyte algae; IRI = 99.98% for ecomorph 1, and IRI = 97.99% for ecomorph 2) and, rarely other items (larvae of water insects - mayflies, chironomids, simulids). The ecomorph 3 had a half-filled gut with dominating periphyton (IRI = 86.3%) with a notable portion of insect larvae (7.62% - predominantly chironomids, also mayflies, and simulids) and macrophytes (5.97%). Ecomorph 4 had fewer filled intestines compared to ecomorph 3 however with strongly dominating periphyton in diet (IRI = 99.49%). The gut of ecomorph 5 (shortest gut) frequently was empty including the largest individual (SL=185 mm). When guts were filled, benthos-associated prey was strongly prevalent (IRI = 99.31%; mayflies and chironomids).

## Mitochondrial data

Both BI and ML analyses of *cytb* revealed monophyly of the *Garra* from the Sore River (Fig. 4A). The closest relative (and ancestor lineage) is from the Barokalu River, a tributary of the Baro River (White Nile drainage). Both Sore and Barokalu rivers share watershed in the Baro system and sampled localities are separated just ca. 50 km by land. Divergence between *Garra* populations from the Sore and Barokalu is low (*p*-distance = 0.0105∓0.0028) and comparable with maximum intra-divergence in the Sore radiation (*p*-distance = 0.0111∓0.0033). Being combined together White Nile lineage is a sister to the large clade of Ethiopian *Garra* from Blue Nile and Lake Tana, Atbara-Nile, Ethiopian Rift Valley, and Omo-Turkana basins.

**Fig. 4.**
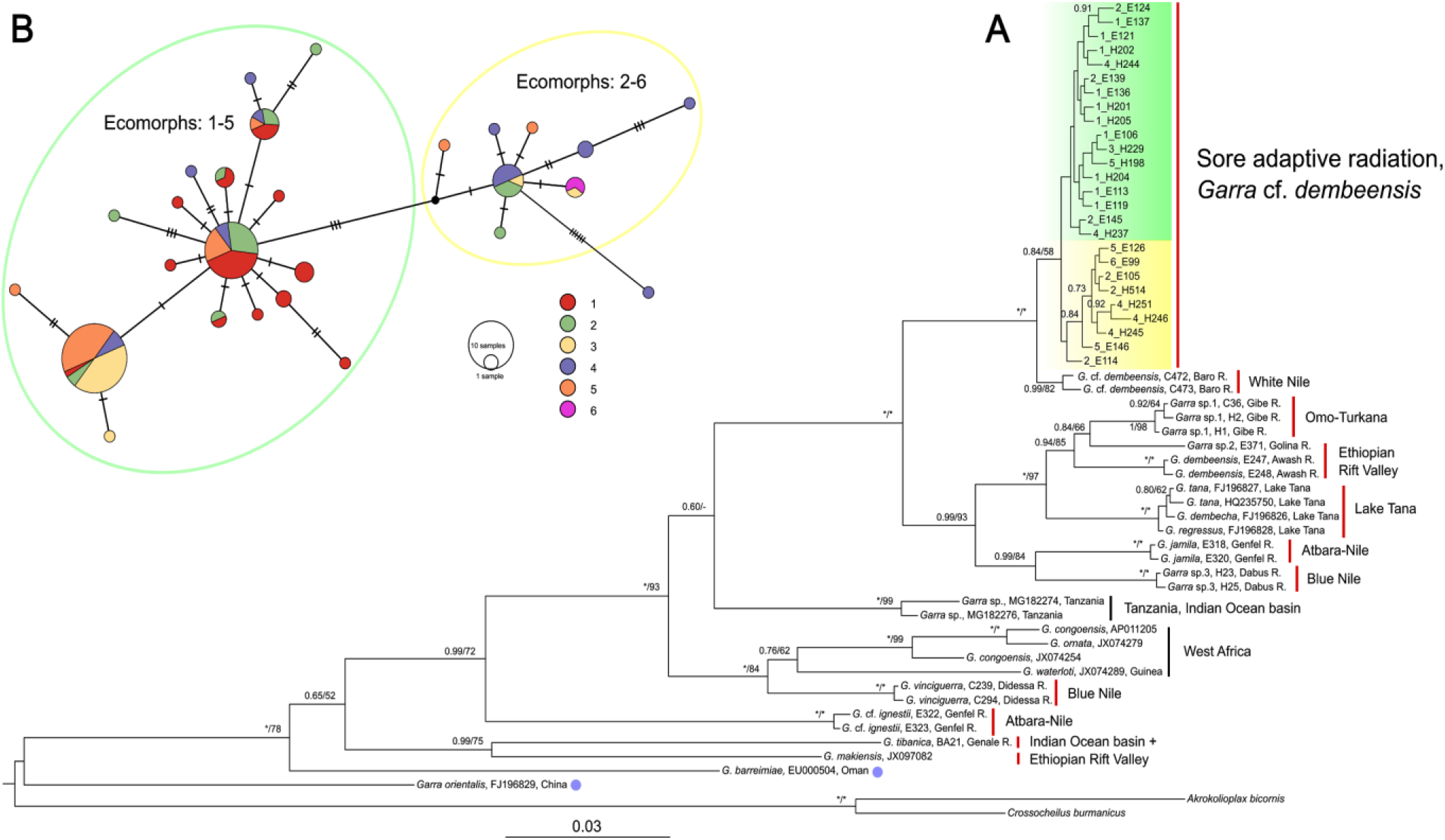
(A) Consensus tree of relationships among the Ethiopian *Garra* from all main drainages based on *cytb* sequences. Bayesian posterior probabilities (before slash) from BI analysis and bootstrap values from ML analysis (after slash) above 0.5/50 are shown; asterisks represent posterior probabilities/bootstrap values of 1/100. Scale bar and branch lengths provide the expected substitutions per site. The green and yellow colors highlight two branches of *Garra* in the Sore River. (B) Median-joining haplotype network of the *Garra* from the Sore River, based on 107 *cytb* sequences (989 bp length). ‘Green’ haplogroup includes ecomorphs 1-5, while ‘yellow’ haplogroup includes ecomorphs 2-6. Black dots represent hypothetical intermediate haplotypes.

At the same time, our phylogenetic analyses revealed that Ethiopian *Garra* are non-monophyletic (Fig. 4A). Some lineages are of more ancient origin and closer to Asian lineages (*G. tibanica* from Indian Ocean basin) or to lineages from West Africa (e.g. *G. vinciguerra* from Blue Nile basin). Matrilineal tree of Ethiopian *Garra* includes up to 12 lineages. Taking into account some species cluster together in one lineage like three species from Lake Tana or that some species were unavailable, we conclude cladogenesis of *Garra* in Ethiopia Highlands has been more diversified than considered previously (Stiassney & Getahun, 2007).

The Sore lineage is composed of two sub-lineages or haplogroups highlighted by yellow and green (Fig. 4A-B). Haplotype net constructed on 107 *cytb* sequences confirms presence of two main haplogroups. The core haplotypes of these haplogroups are separated by 5 substitutions. Four of six ecomorphs (2, 3, 4, and 5) share both haplogroups. The ‘green’ haplogroup is prevalent in number of haplotypes (18), and number of individuals (88), and found in five ecomorphs. Ecomorph 1 is presented exclusively in this haplogroup. In contrast, the ‘yellow’ haplogroup (Fig. 4B) is smaller, with only different 9 haplotypes found in 19 individuals (= 17.7 % of the individuals analyzed). One individual of ecomorph 4 is rather distant (6 substitutions) from the core haplotype of this haplogroup. ‘Yellow’ haplogroup consists of five ecomorphs as well. However, ecomorph 4 is much more frequently represented in this haplogroup (42 % of all individuals) compared to ‘green’ one (6.97 %).

## RAD-seq data

Raw reads statistics is given in Supplementary File S1.

### Nuclear phylogeny

The phylogeny of Ethiopian *Garra* based on a concatenated set of RAD-loci sequences (23,365 partitions and 3,075,180 total sites with 0% missing data) is generally similar to that based on mtDNA data (Fig. 4) but it has more strongly supported nodes, as it is based on many more variable sites (Fig. 5A). Sympatric ecomorphs clustered together and form monophyletic lineages, sister to the population from the same riverine basin - Baro drainage in White Nile system (Fig. 5A-B). Closest relative to *Garra* from White Nile system is *Garra* lineage in the *G. dembeensis* complex from neighbor drainage - Omo-Turkana system. The *G. vinciguerrae* from the Blue Nile (which recorded in Ethiopia for the first time in the current study) is ancestor lineage for both White Nile and Omo-Turkana lineages. The most divergent lineages, *G. makiensis* and *G. tibanica*, are from Ethiopian Rift Valley and Indian Ocean basins, respectively.

**Fig. 5.**
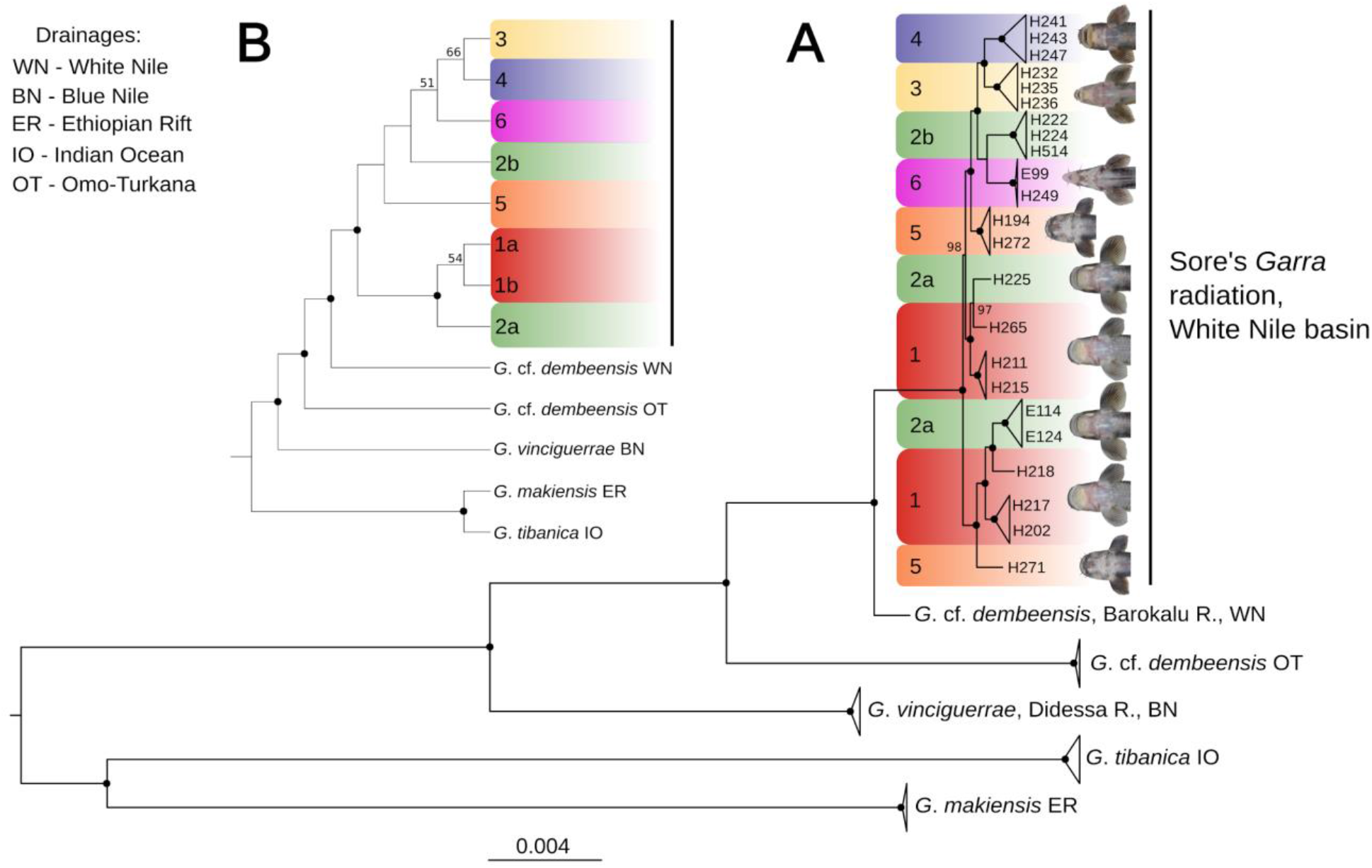
(A) ML phylogenetic tree of Ethiopian *Garra* based on RAD-loci sequences - 23,365 loci; 3,075,180 bp and (B) SVDQ species tree. Each locus was treated as a separate partition with GTR+G substitution model and heterozygous sites within each individual encoded using IUPAC notation. Black dots designate 100% bootstrap support, and only values above 50% are given.

Compared to mitochondrial data, the nuclear phylogenomic tree shows much better segregation of *Garra* ecomorphs from the Sore River (Fig. 5A). Ecomorphs 3, 4, and 6 form monophyletic clusters, while other ecomorphs are divided into two (nos. 1 and 5) or even three (no. 2) clusters. We assign two distantly located branches of both ecomorph 1 (generalized) as 1a/1b as well as ecomorph 2 (stream-lined) as 2a/2b according to population genomics analyses done below (Fig. 6-8). Ecomorphs 1 and 2 from one hand, and other ecomorphs from another hand form two clusters within Sore River adaptive radiation according to SVDQ species tree (Fig. 5B). Ecomorphs 3 (narrow-mouth), 4 (wide-mouth), and 6 (thick-lipped) are most recently diverged branches according to SVDQ-tree but the nodes are weakly supported (Fig. 5B).

**Fig. 6.**
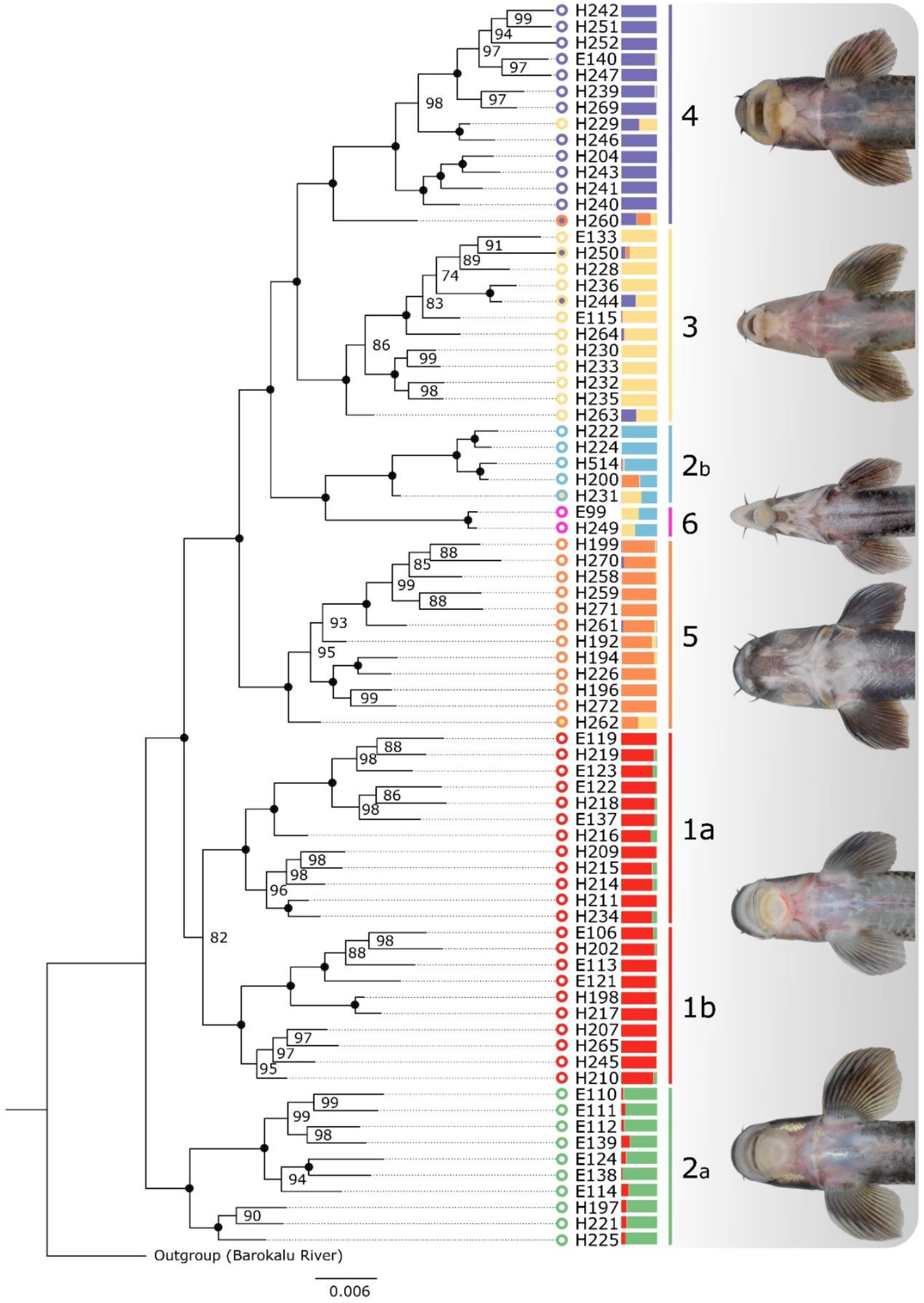
ML phylogeny of sympatric *Garra* ecomorphs from the Sore River based on concatenated RAD-loci sequences (7,370 loci; 969,450 bp). Each locus was treated as a separate partition with GTR+G substitution model. Heterozygous sites within each individual encoded using IUPAC notation. The individual samples are colored based on the color scheme of Fig. 4 and intermediate (putative hybrids) phenotypes are depicted in another color. The genetic clusters proportions inferred by *rmaverick* analysis are shown to the right of sample numbers. Black points designate 100% bootstrap support.

**Fig. 7.**
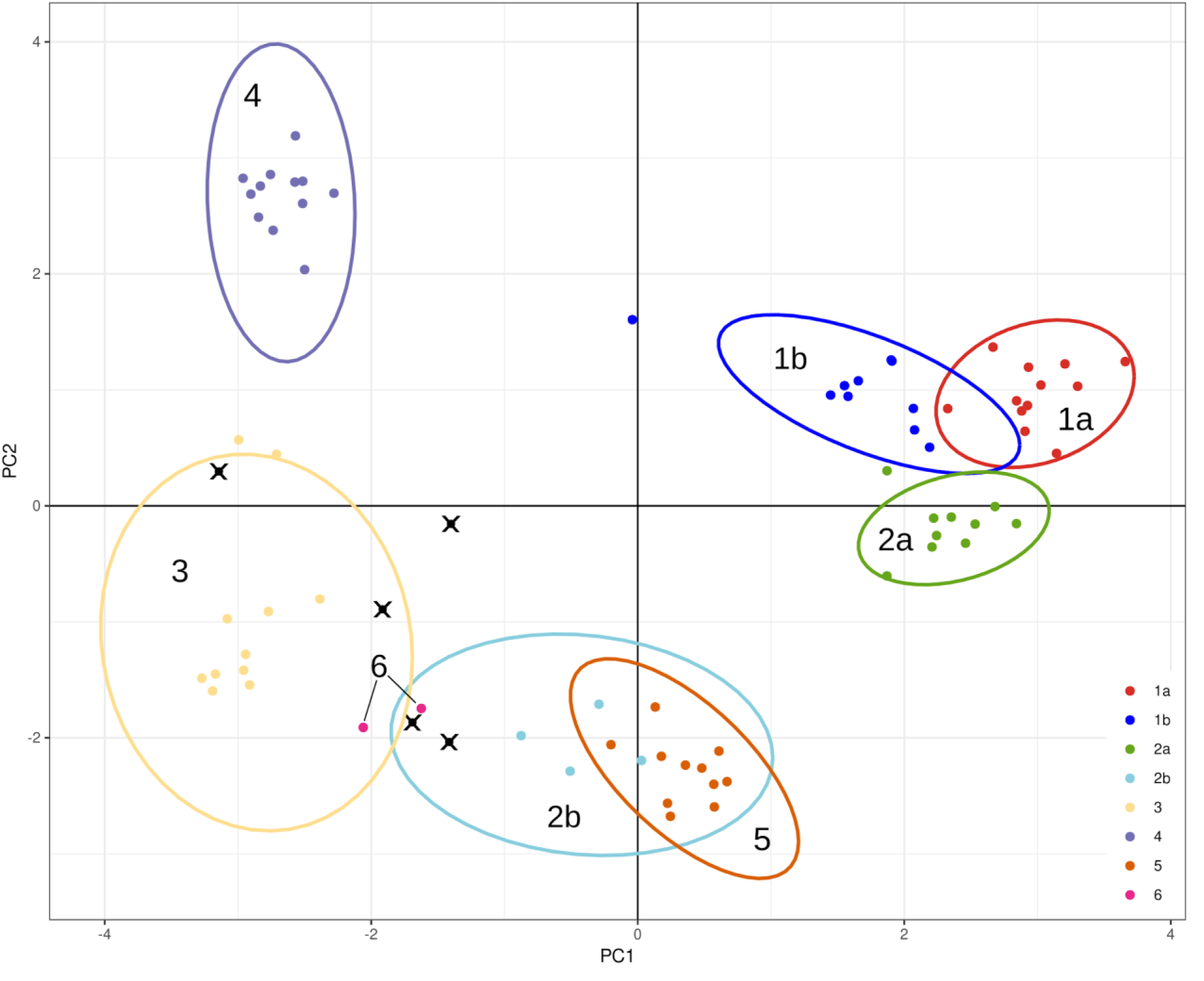
Principal Component Analysis (PCA) based on 679 nuclear SNPs of sympatric *Garra* ecomorphs from the Sore River. Points (individuals) and 95% confidence ellipses are colored by phenotype/genetic cluster. Crosses assign intermediate phenotypes.

**Fig. 8.**
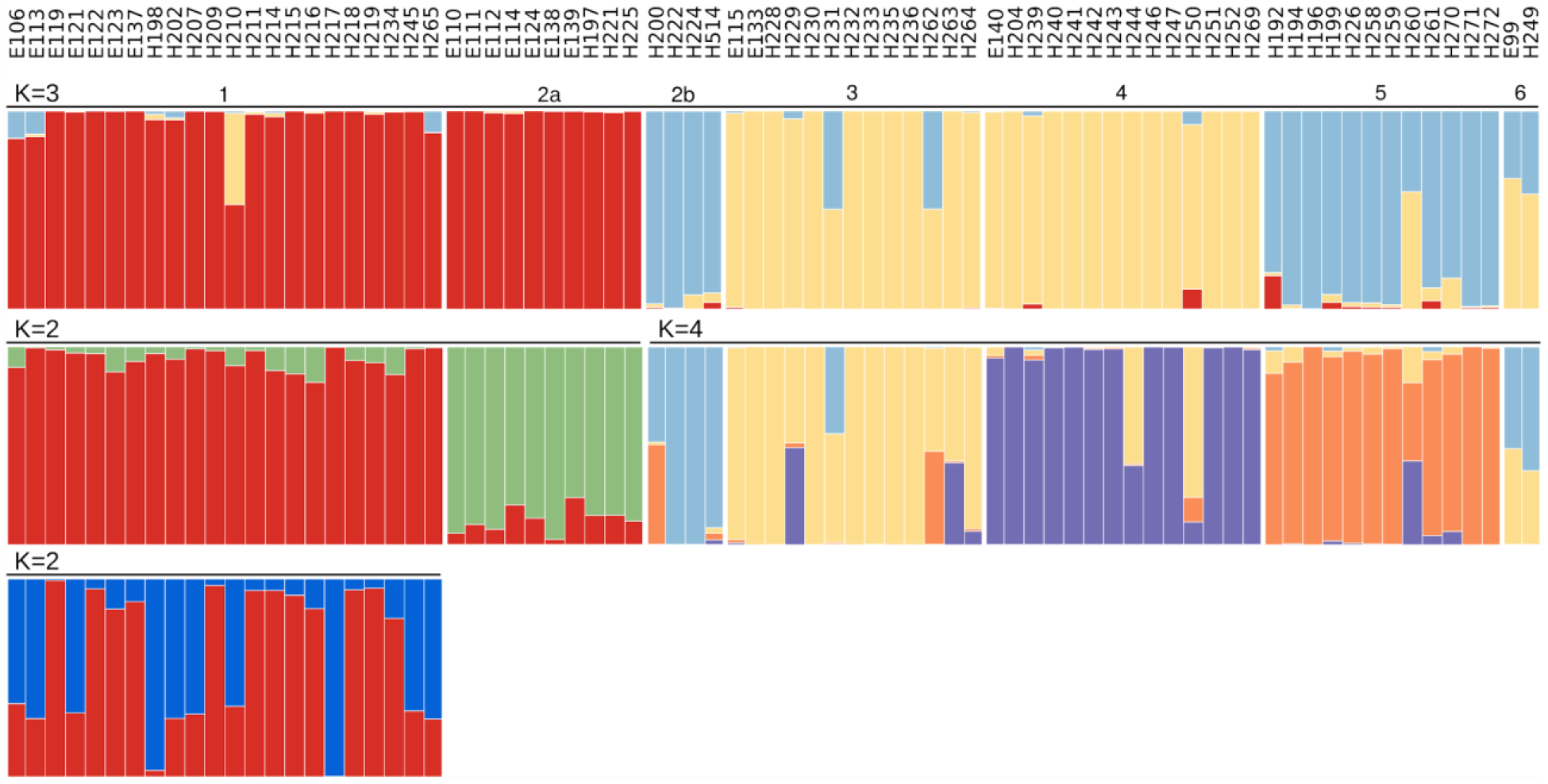
Hierarchical *rmaverick* results for sympatric ecomorphs of *Garra* from the Sore River, based on 679 nuclear SNPs. Each column of the barplot shows individual assignments to one of the inferred genetic clusters. Independent runs of *rmaverick* are indicated by a solid black line above a plot, along with an inferred value of *K*.

Relationships among the Sore River sympatric ecomorphs based on analysis of all samples and full RAD-loci sequences (> 7000 loci and > 0.96 Mbp length sequences) are presented in Fig. 6. The ML analysis highly support the monophyly of each ecomorph except for ecomorph 2. The most basal lineage is ecomorph 2, which in turn, is paraphyletic, suggesting, possibly, that there is another 7^th^ cryptic species that we could not distinguish phenotypically. Four individuals along with one individual of intermediate phenotype represent another lineage that we call 2b (Fig. 6). Lineage 2a is sister all other ecomorphs that are divided for two subclades - one includes only ecomorph 1 individuals (which, in turn is subdivided into what we call - 1a-1b), while another includes all other ecomorphs - 3, 4, 5, 6, and above mentioned 2b. That latter lineage is composed of lineages, each containing samples of particular ecomorphs except for several samples which were intermediate in their phenotypes (Fig. 6). Ecomorph 6 (thick-lipped mouth) is resolved as sister to the 2b lineage albeit with an apparent rather deep last common ancestor. Generally, the placement of clade 2a as sister to all other *Garra* from the Sore River, that is characterized by a well-developed gular disc (type C), might suggest that this an ancestral condition of this radiation.

### Population genomics

Principle component analyses of the 679 nuclear SNPs of sympatric ecomorphs revealed several well-defined clusters that correspond to the phenotypic differentiation (Fig. 7). Ecomorph 1 (composed of two genetic sub-clusters 1a-1b), genetic cluster 2a, ecomorphs 3 and 4 are not overlapping, while clusters of 2b and ecomorph 5 broadly overlap. Thick-lipped ecomorph (6) interestingly (although it is difficult to place since we only found two individuals that we could include in this study) could not be identified by PCA as a distinct cluster.

The analysis of population structure with admixture revealed an optimum of three genomic clusters that correspond to the i) ecomorph 1 + 2a lineage, ii) ecomorphs 3 + 4, and iii) ecomorph 5 + 2b lineage (Fig. 8, Upper row, K3). Ecomorph 6 is characterized by admixture of two clusters from ecomorphs 3 and 4.

Subsequent analysis of each cluster (=lineage) revealed hierarchical subdivision. Thus ecomorph 1 and genetic lineage 2a each are also identified as cluster in the admixture analysis (Fig. 8 middle row, K=2). Although ecomorphs 3, 4, 5, and lineage 2b are supported as independent evolutionary units based on several types of genetic analyses, few individuals in all of these show signs of historical gene flow based on the admixture analysis (Fig. 8). While the two individuals from ecomorph 6 in our study seem most clearly be composed of genetic contributions by ecomorphs 3 (36.8-47.5%) and genetic lineage 2b (51.3-62.3%), possibly supporting a hybrid origin hypothesis. Interestingly, one more individual with combination of the same genomic clusters but with the opposite ratio (54.0% from ecomorph 3 and 43.9 % from lineage 2b) had no thick-lipped features (the main phenotypic diagnostic feature for ecomorph 6) and was phenotypically assigned to ecomorph 3 (Fig. 8). One more level of population subdivision was detected in ecomorph 1 (Fig. 8) with two genomic clusters (lineages 1a and 1b) of high degree of admixture. It suggests heterogeneous genomic structure of the generalized ecomorph as a result of secondary contact.

All Reich F_ST_ pairwise comparisons were statistically significant with values ranging from 0.10 (lineages 1a vs. 1b) to 0.46 (ecomorphs 2b vs. 6) (Fig. 9). The ecomorph 6 F_ST_ values were the highest (0.39-0.46).

**Fig. 9.**
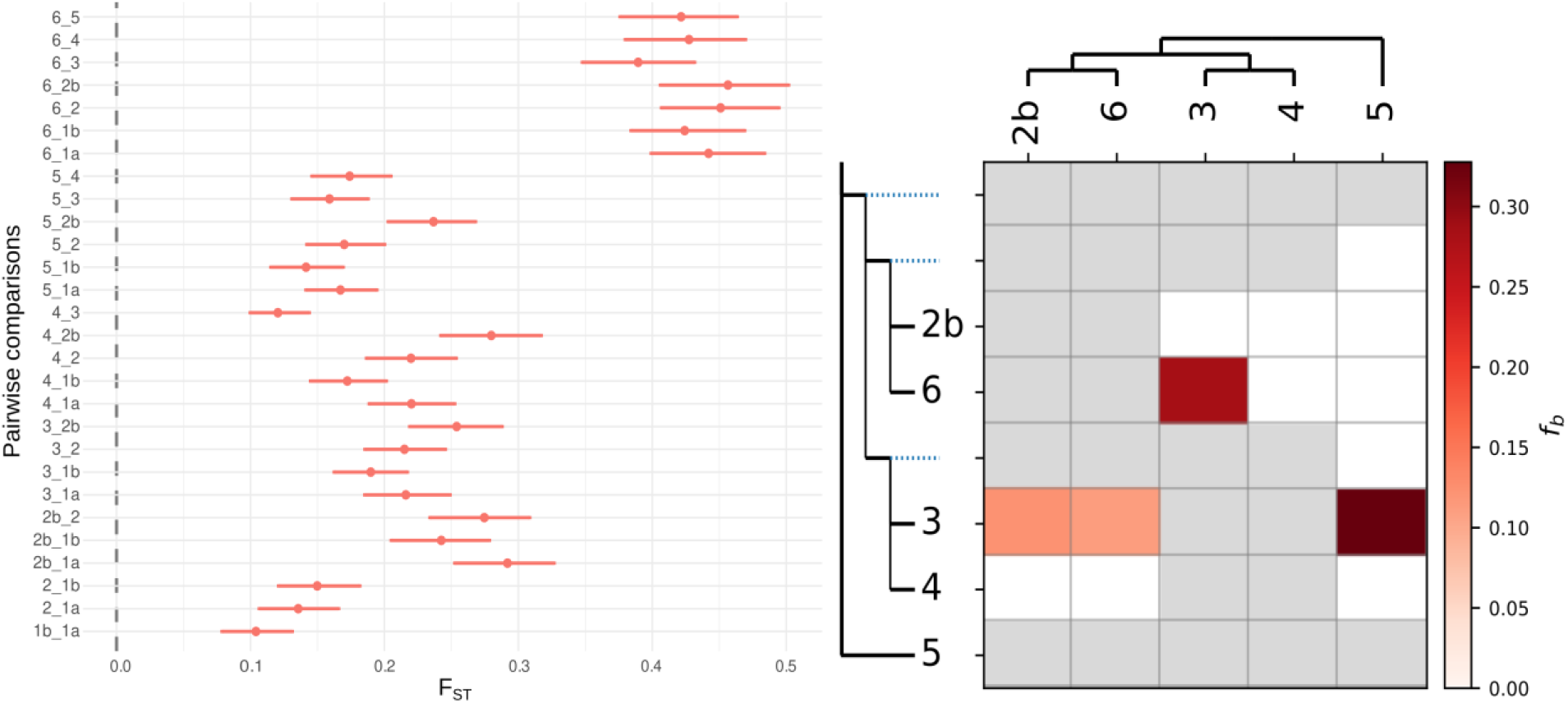
Left - pairwise Reich F_ST_ values (points) with their respective 95% confidence intervals (horizontal lines) for *Garra* genetic lineages from the Sore River based on 679 SNPs. Right - heat map of ƒ-branch metric for selected ecomorphs/lineages of the *Garra* Sore radiation. The used guide tree is shown along the x and y axes (in ‘laddered’ form along the y axis). The matrix shows the inferred ƒ-branch metric, reflecting excess allele sharing between the branch of the ‘laddered’ tree on the y axis (relative to its sister branch) and the branches defined on the x axis.

As the *rmaverick* analysis suggested a notable level of admixture between lineage 2b and ecomorphs 3, 4, and 6 (Fig X), which form a single monophyletic cluster in our phylogenomic analysis (Fig 8), we performed a number of tests to distinguish between gene flow (introgression) and incomplete lineage sorting (ILS). The obtained D statistic was positive and significant for a number of comparisons (Table 3.). Visualization of ƒ-branch metric (which is based on ƒ_4_-ratio results) highlighting introgression between ecomorphs/genetic lineages 2b and 3, 6 and 3, 5 and 3 (Fig 9).

**Table 3.** Results of Patterson’s D statistic (ABBA-BABA test) and ƒ4-ratio test on selected genetic clusters of *Garra* from the Sore River.

| P1 | P2 | P3 | D statistic | Z-score | p-value | f4-ratio | BBAA | ABBA | BABA |
| --- | --- | --- | --- | --- | --- | --- | --- | --- | --- |
| 4 | 3 | 6 | 0.1176 | 5.3829 | **<0.0001** | 0.1128 | 227.5 | 235.0 | 185.5 |
| 2b | 3 | 5 | 0.0650 | 3.1078 | **0.0009** | 0.4226 | 253.5 | 246.5 | 216.4 |
| 2b | 6 | 3 | 0.0646 | 2.3475 | **0.0095** | 0.2854 | 215.6 | 217.3 | 190.9 |
| 4 | 3 | 2b | 0.0624 | 3.8143 | **<0.0001** | 0.1237 | 264.6 | 241.4 | 213.0 |
| 4 | 3 | 5 | 0.0492 | 3.6742 | **0.0001** | 0.3277 | 276.2 | 247.4 | 224.2 |
| 2b | 6 | 5 | 0.0327 | 1.4755 | 0.0700 | 0.2051 | 248.6 | 203.4 | 190.5 |
| 4 | 6 | 5 | 0.0304 | 1.5315 | 0.0628 | 0.2330 | 224.5 | 226.5 | 213.2 |
| 6 | 3 | 5 | 0.0199 | 0.9380 | 0.1741 | 0.1641 | 244.2 | 204.7 | 196.8 |
| 2b | 4 | 5 | 0.0178 | 1.0774 | 0.1406 | 0.1134 | 245.9 | 246.3 | 237.7 |
| 2b | 6 | 4 | 0.0040 | 0.1592 | 0.4368 | 0.0151 | 244.6 | 197.8 | 196.3 |

The eighth genetic clusters possess from three (ecomorph 6) to 38 private alleles (ecomorph 4) (Table 4). The ecomorph 6 has also the lowest heterozygosity (Ho = 0.00058) as well as nucleotide diversity (Pi = 0.00054) compared to all other ecomorphs (Ho = 0.00104-0.00128; Pi = 0.00121-0.00091) (Table 4).

**Table 4.** Summary of the ecomorphs’ genetic diversity indices averaged over 89 070 loci (both variant and fixed).

| Ecomorphs * | No. of private alleles, Np | No. of polymorphic loci, % | Heterozygosity | | Coefficient of inbreeding (Fis) ± SE | Nucleotide diversity (Pi) ± SE |
| --- | --- | --- | --- | --- | --- | --- |
| | | | Observed (Ho) ± SE | Expected (He) ± SE | | |
| 1a | 19 | 0.42 | 0.00128±0.00008 | 0.00116±0.00007 | -0.00014±0.0015 | 0.00121±0.00007 |
| 1b | 18 | 0.40 | 0.00128±0.00008 | 0.00113±0.00007 | -0.00019±0.0011 | 0.00119±0.00007 |
| 2a | 27 | 0.41 | 0.00124±0.00008 | 0.00114±0.00007 | -0.00007±0.0012 | 0.00120±0.00007 |
| 2b | 9 | 0.24 | 0.00104±0.00008 | 0.00079±0.00006 | -0.00023±0.0012 | 0.00091±0.00007 |
| 3 | 20 | 0.43 | 0.00127±0.00008 | 0.00107±0.00006 | -0.00037±0.0013 | 0.00111±0.00007 |
| 4 | 38 | 0.43 | 0.00109±0.00007 | 0.001±0.00006 | -0.00008±0.0015 | 0.00104±0.00006 |
| 5 | 33 | 0.44 | 0.00126±0.00008 | 0.00115±0.00007 | -0.00011±0.0019 | 0.00120±0.00007 |
| 6 | 3 | 0.10 | 0.00058±0.00007 | 0.0004±0.0000 | -0.00006±0.0004 | 0.00054±0.00006 |

\* - letters 'a' and 'b' assign genetic lineages within ecomorphs 1 and 2.

## Discussion

Our study provides genetic support for the hypothesis of the evolution of an adaptive radiation in a riverine environment. By analyzing trophic features and sucking disc variation, as well as trophic ecology, we show morpho-ecological diversification of the cyprinid fish *Garra dembeensis* into six distinct ecomorphs. First, diversification of two novel phenotypes (thick-lipped and predatory) in the Sore River has evolved rapidly, an event that can be classified as burst of speciation sensu Givnish (2015). Second, adaptive radiation resulted in the origin of several highly specialized lineages of algae scrapers, i.e. specialized ancestor adaptively radiates giving rise to eco-morphological diverse lineages, that seem to be not only ecologically, but also reproductively isolated from each other and can be considered the new species.

### Eco-morphological diversification and adaptive radiation of Garra

The genus *Garra* is currently comprised of more than 160 species (Fricke et al., 2021; Yang et al., 2012). Only 23 of which occur in Africa (Moritz et al., 2019). So far, 13 described species were reported from Ethiopia (Golubtsov et al., 2002; Stiassny & Getahun, 2007). In this study, we discovered six additional distinct ecomorphs that originated through adaptive radiation in the Sore River, and thus might warrant the description of five-six new African *Garra* species.

The ecomorphs of the Sore’s *Garra* are exceptionally diverse in trophic and sucking disc morphology. Two novel phenotypes for the whole genus *Garra* – ‘thick-lipped’ and ‘predatory’ - have superficial similarities to Lake Tana large barbs species/morphotypes, e.g., thick-lipped barb *L. negdia* (Rüppell, 1836) and predatory *L. gorguari* (Rüppell, 1836) (Nagelkerke & Sibbing, 1997). This high degree of variation in the sucking disc in Sore’s *Garra* can be observed - from well-developed disc with free posterior margin to complete absence. Such a degree of morphological diversity concentrated in one riverine spot of Ethiopian Highlands would seem to satisfy the requirements of a diversification burst (sensu Givnish, 2015).

Divergent feeding-related morphology and gut content analysis suggest trophic specialization of *Garra* sympatric forms. This is consistent with other cases of adaptive radiation among Ethiopian cyprinids, where trophic resource partitioning promoted diversification - *Labeobarbus* spp. in Lake Tana (Sibbing, Nagelkerke, Stet, & Osse, 1998) as well as in the Genale River (Levin et al., 2019). The most common foraging strategy among *Garra* is scraping of periphyton from stones and rocks (Hamidan, Jackson, & Britton, 2016; Matthes, 1963). This is predominant in Sore’s *Garra* ecomorphs 1 and 2 that have long gut (4-5 times longer than body length) filled with periphyton and detritus. The ecomorphs 1 and 2 are divergent mainly in body shape. The latter has streamlined appearance and probably is adapted for life in more rapid flowing water. Ecomorph 3 has shorter gut length (ca. 2-times longer than body length) and a mixed diet with significant additions of benthic invertebrates. Ecomorph 5 has an extremely short gut, whose length is as long as the fish body. Short gut is a strong marker for predatory/piscivory feeding strategy in fishes, including cyprinids (Nagelkerke, 1997; Sibbing et al., 1998; Wagner, McIntyre, Buels, Gilbert, & Michel, 2009, Zandoná, Auer, Kilham, & Reznick, 2015). Predatory *Garra* from the Sore River have 4-5-times shorter gut length than congeneric periphyton feeders and twice shorter gut than that of piscivory large-mouthed ecomorph of *Labeobarbus* from the Genale River, Ethiopia (Levin et al., 2019). We found an empty gut in many individuals of ecomorph 5, while small-sized fishes had gut filled with insects. Ecomorph 4 has a rather long intestine and predominantly periphyton in diet, but it is characterized by distinctly divergent mouth phenotype compared to ecomorphs 1 and 2 (Fig. 3). The gut of thick-lipped phenotype (ecomorph 6) was not analyzed because of the extreme rarity of samples. Hypertrophied lips (or ‘rubber lips’) of fishes is an adaptation to foraging on benthos hidden between rock crevices on pebble and rock fragments via increased sucking power by sealing cracks and grooves (Baumgarten, Machado-Schiaffino, Henning, & Meyer, 2015; Machado-Schiaffino, Henning, & Meyer, 2014; Matthes, 1963; Ribbink, Marsh, Marsh, & Sharp, 1983). This phenotype is widely distributed among other cyprinid fish, the *Labeobarbus* spp., inhabiting lakes and rivers of Ethiopian Highlands (Mina, Mironovsky, & Dgebuadze, 1996; Mironovsky, Mina, & Dgebuadze, 2019; Nagelkerke, Sibbing, van den Boogaart, Lammens, & Osse, 1994) including the Sore River (Levin et al., 2020), but it was never detected among *Garra* species. Our study shows that the thick-lipped mouth phenotype represents an evolutionary novelty within the *Garra* lineage that most probably resulted from hybridization events between ecomorphs 2 (lineage 2b) and 3 because its genome had an admixture from these genetic lineages. Hybridogenic origin of the *Garra*’s thick-lipped phenotype may corroborate results of recent experimental study demonstrating the importance of hybridization in generating of functional novelty of ecological relevance in relation to trophic resources unavailable for parental species in cichlids (Selz & Seehausen, 2019). The origin of novel thick-lipped phenotype in the genus *Garra* is of particular interest in light of knowledge of non-hybrid origin of hypertrophied lips from ancestors with normally developed lips in cichlid fishes (Baumgarten et al., 2015; Machado-Schiaffino et al., 2017). Interestingly, there might only be a single locus involved in producing the hypertrophied cichlid phenotype (Kautt et al., 2020), the genomic basis of the lip phenotypes in *Garra* remains unknown.

Another novel phenotype for *Garra* detected in the Sore River is the “predatory” niche. A conspicuously piscivory trophic strategy is rare among Cypriniformes, presumably because they have a toothless jaw. Nevertheless, this feeding strategy is quite common among cyprinid fishes inhabiting water bodies of Ethiopian Highlands. For example, seven of the total 15 endemic *Labeobarbus* spp. found in Lake Tana are predatory on fish (Nagelkerke et al., 1994; Sibbing et al., 1998); that evolved multiple times among riverine populations of the genus *Labeobarbus* (Levin et al., 2020).

To our knowledge, only one sympatric diversification has previously suggested for *Garra* – the intralacustrine complex including three species inhabited Lake Tana in Ethiopia (Geremew, 2007; Stiassny & Getahun, 2007). This diversification resulted in divergent phenotypes (gular discs varies from well-developed to reduced size) and ecology (one form is pelagic - *G. tana*) and can be considered as a recent speciation as suggested by the absence of mtDNA divergence among these species (Tang, Getahun, & Liu, 2009). Unfortunately, little is known about morpho-ecological and genetic diversity of this Lake Tana radiation. Sympatric divergence was also recently proposed as the most likely mechanisms for the origin of two blind *Garra* species, *G. typhlops* and *G. lorestanensis*, inhabited the same cave in Zagros Mountains, Iran (Segherloo et al., 2018).

### Possible scenarios of evolution of Garra’s adaptive radiation in the Sore River

Both mtDNA and genome-wide SNPs data support monophyly of the Sore’s *Garra* as well as their recent speciation based on low genetic divergence between the nearest ancestor and Sore River’s ecomorphs. The closest relative and ancestor of the Sore River diversification inhabits the same subbasin of the White Nile in Ethiopia, therefore suggesting an intra-basin diversification of *Garra* there. On the one hand, mtDNA data might have failed to distinguish sympatric ecomorphs because of high level of shared genetic diversity caused by ILS and introgression, this latter highlighted by D-statistic calculated with the genome-wide nuclear data. On the other hand, the SNP data support a reproductive isolation among closely-related ecomorphs despite few individuals having intermediate phenotypes and genetic admixture. Hybrid origin of intermediate phenotypes might suggest that reproductive isolation barriers are not complete yet.

Patterns of haplotype net (numerous haplotypes occurring in the same phenotypes) as well as SNP data (presence of more genetic clusters than phenotypes) could also suggest secondary contact of local sub-isolated populations. The riverine net of Ethiopian Highlands was significantly influenced by several episodes of dramatic volcanism and tectonism until the Quaternary (Ferguson et al., 2010; Hutchison et al., 2016; Prave et al., 2016). Thus, riverine net fragmentation, isolation or sub-isolation of some riverine parts, and captures of headwaters is a likely scenario given the geological history of Ethiopian Highlands (Mège, Purcell, Pochat, & Guidat, 2015), also supported by genetic studies on other Ethiopian fishes (Levin et al., 2019; 2020). Concerning the Sore River, while waterfalls and rapids are rather frequent, no geological data that support its connection to other basins are known. In our view, the most reliable evolutionary scenario for the origin of the riverine adaptive radiation in the *Garra* species group draws upon a combination of allopatric and sympatric stages of speciation with hybridization and admixture. A comparable evolutionary history was detected in the *Labeobarbus* adaptive radiation in the Genale River (Ethiopia), which is part of the extended ancient riverine net in Juba-Wabe-Shebelle drainage (Levin et al., 2019).

Speciation with gene flow was detected in several studies (e.g. Feder, Egan, & Nosil, 2012; Fruciano, Franchini, Raffini, Fan, & Meyer, 2016; Kautt, Machado-Schiaffino, & Meyer, 2016; Kautt et al., 2018; Kautt et al., 2020; Machado-Shiaffino et al., 2017; Malinsky et al., 2018; Puebla, 2009; Rougeux, Bernatchez, & Gagnaire, 2017; Schwarzer et al., 2011; Smadja & Butlin, 2011; Zheng & Ge, 2010). Notably, it has been shown as genetic admixture between divergent populations/lineages may be a key factor in promoting rapid ecological speciation (Jacobs et al., 2020; Kautt et al., 2016; Kautt et al., 2020; Martin et al., 2015; Marques, Meier, & Seehausen, 2019). Moreover, ancient hybridization is widely considered one of the most important factors driving the spectacular cichlid adaptive radiations in the Great African Lakes (Irissari et al., 2018; Meier et al., 2017; Verheyen, Salzburger, Snoeks, & Meyer, 2003). Seemingly, ancient introgressive hybridization could be a trigger for small-scaled repeated adaptive radiations among the Arctic charrs *Salvelinus* (Lecaudey et al., 2018). Furthermore, hybridization is the main mechanism generating polyploid lineages in fishes (tetraploid, hexaploid etc. - Braasch & Postlethwait, 2012), whose complex genomes constitute the raw material for the rapid origin of sympatric forms (e.g. *Schizothorax* in Central Asia - Berg, 1914; Burnashev, 1952; Terashima, 1984; *Labeobarbus* in Africa - Levin et al., 2020; Mina et al., 1996; Nagelkerke et al., 1994; Vreven, Musschoot, Snoeks, & Schliewen, 2016). Nevertheless, all described *Garra*, including the Ethiopian species, have diploid genomes (Krysanov & Golubtsov, 1993).

### Adaptive radiation in riverine environment

Most adaptive radiations of fishes were reported from the lacustrine environment (e.g., Fryer & Iles 1972; Seehausen & Wagner, 2014). However, increasing evidence suggest that adaptive radiation can take place in other aquatic environments (e.g., marine, riverine) (Burress et al., 2018; Dimmick et al., 2001; Feulner, Kirschbaum, & Tiedemann, 2008; Levin et al., 2019; 2020; Melnik et al., 2020; Matchiner, Hanel, & Salzburger, 2011; Piálek et al., 2012; Puebla, 2009; Whiteley, 2007). Several other cases of potential riverine adaptive radiations that includes ≥ 3 sympatric ecomorphs exist, although they were not been tested with genetic methods yet - for instance, snow trout from Central Asia (Berg, 1914; Burnashev, 1952), barbs *Poropuntius* and *Neolissochilus* from Southeastern Asia (Roberts, 1998; Roberts & Khaironizam, 2008). Among cichlids, one of the first riverine adaptive radiations examined genetically were from Southern Africa (Joyce et al., 2005). However, the authors of this study suggested that the adaptive radiation occurred in the lacustrine environment in the palaeo lake Makgadikgadi that dried up in the Holocene (Joyce et al., 2005). Other cichlid adaptive radiations from the rivers of Western Africa (Schwarzer et al., 2011), Southern America (Burress et al., 2018; Piálek et al., 2012;) as well as four independently evolved riverine radiations of labeobarbs from East Africa (Levin et al., 2020), have instead took place in riverine drainages without known lacustrine conditions in the past.

The *Garra* lineage is adapted to fast and torrent waters. It possesses a morphological novelty - gular sucking disc - used to cling on the bottom of swift waters. This novelty allowed *Garra* to be distributed widely in highlands and montane zones from Southeastern China to Western Africa. Only a few species were found in the lacustrine environment (Lake Tana – Stiassny & Getahun, 2007) or in caves (e.g. Banister, 1984; Coad, 1996; Kruckenhauser, Haring, Seemann, & Sattmann, 2011; Mousavi-Sabet & Eagderi, 2016), indicating their potential to adapt to steady waters.

Despite the riverine network is generally considered more open to gene flow compared to landlocked water bodies, mountain and highland are an exception to this rule. The Ethiopian Highlands are a volcanic massif of flood and shield volcano basalts 0.5–3.0 km thick that form spectacular trap topography (1500–4500 m) flanking the Main Ethiopian Rift (Prave et al., 2016). The geological history of the Ethiopian Highlands was tectonically very dynamic and rich in volcanic episodes from Oligocene to Pleistocene time with very recent episodes (Prave et al., 2016). The volcanic activity has been severe enough to deleteriously affect the biota and cause major disruptions in ecosystems. This hypothesis found support in the inferred evolutionary history of the *Labeobarbus* in East Africa. The earliest fossil records of *Labeobarbus* were found in the Ethiopian Rift Valley and dated back to the late-Miocene (Stewart & Murray, 2017), but most of the Ethiopian lineages are younger (Pleistocene origin) (Beshera, Harris, & Mayden, 2016; de Graaf, Megens, Samallo, & Sibbing, 2010; Levin et al., 2020). The tectonic activity of the region could have favored local isolation via the formation of waterfalls (e.g., 33 kya the Blue Nile basaltic blockade formed Tis-Isat waterfall - Prave et al., 2016) or river net fragmentation (Juba-Wabe-Shebelle drainage Mège et al., 2015) along with climatic oscillations resulted to disconnection of water bodies during aridization (Benvenutti et al., 2002). Periodically, it resulted in vacant habitats and ecological opportunity (reviewed by Stroud & Losos 2018) for new species to exploit similar to islands or crater lakes (Burress et al., 2018).

The *Garra*’s diversification burst in the Sore River was detected in the riverine segment at an altitude range of 1310-1550 m asl, that is within the range of four riverine diversifications of the *Labeobarbus* detected throughout Ethiopian Highlands: 1050-1550 m (Levin et al., 2020). Despite the generally broader elevation gradient (175-2000 m asl - Levin et al., 2020) of the *Labeobarbus* species complex, the diversification bursts were only detected in mid-upper reaches. We believe that a combination of two factors might explain this observation: i) fauna in mid-upper reaches is poorer compared to lower reaches, where a more diversified fauna might have already filled the available ecological niches necessary for an adaptive radiation to unfold; ii) the biotopes are more diverse compared to the most upper reach, that means vacant niches are available.

Five endemic, and one introduced non-*Garra* species were recorded in the Sore River in the study area (data of this study). This is an extremely low number compared to more than 110 fish species (Golubtsov & Darkov, 2008, and our data) recorded in the Baro River at Gambella at 440m altitude (our data) to which the drainage of the Sore River belongs with a distance of ∼150km between compared localities. The segment of the Sore River where *Garra’*s diversification was detected is rather rich in biotope complexity - pools are alternating pools slow currents, rift areas and rapids (Fig. S6). The depauperated fauna was suggested to provide the ecological opportunities for riverine adaptive radiations similar to the in Southeastern cyprinids of the genus *Poropuntius* (Roberts, 1998) and South America cichlids of the *Crenicichla* due to relaxed competition and vacant niches might have provided ecological opportunities for sympatric speciation by trophic specializations (Burress et al., 2018).

We discovered six new species within the genus *Garra* in the Sore River. Given that the same riverine segment is home for another riverine diversification of fishes represented by four phenotypically diverged ecomorphs of the genus *Labeobarbus* (Levin et al., 2020), we consider the Sore River to being a hot-spot of riverine diversification in the Ethiopian Highlands that requires conservation management. The Ethiopian Highlands are home for several young fish radiations - a large lacustrine diversification among cyprinids (15 species/morphotypes - Mina et al., 1996; Nagelkerke et al., 1994; Nagelkerke et al., 2015) as well as small-sized diversifications of *Garra* (three species – Stiassny & Getahun, 2007) and *Enteromius* (two species - de Graaf, Megens, Samallo, & Sibbing, 2007; Dejen et al., 2002) - all in Lake Tana, and five riverine adaptive radiations of cyprinids each including from four to seven species (Golubtsov, 2010; Golubtsov, Korostelev, & Levin, 2021; Levin et al., 2019; 2020; Mina, Mironovsky, Golubtsov, & Dgebuadze, 1998; current study), highlighting this region’s importance as a hotspot for fish speciation that is in need of additional research on ecological speciation processes.

## Acknowledgements

The study was supported by Russian Science Foundation (grant no. 19-14-00218). We are grateful to all members of Joint Ethiopian-Russian Biological Expedition (JERBE), who participated in our field operations (S.E. Cherenkov, Genanaw Tesfaye, Fekadu Tefera, and I.S. Razgon), and especially to JERBE coordinator Dr. A.A. Darkov for his permanent and invaluable aid. We are thankful to O.N. Artaev for creating a map, S.E. Cherenkov for photographing the fish, A.S. Komarova for data on the gut content as well as to M.V. Mina for critical notes on the first variant of manuscript.

## Author contributions

BL, ES, PF, NM, AG, and AM designed and contributed to the original concept of the studies. BL and AG collected most of the specimens and related data, BL and NM obtained mtDNA data and prepared DNA libraries for ddRAD, BL conducted morphologic analyses, ES conducted the most of bioinformatics, and BL, ES, PF, and AM finalized the manuscript. All authors participated in project design, and read and approved the final manuscript.

## Data availability statement

Morphologic data (body proportions and gut lengths), mtDNA subsets (cytochrome *b*), and genotyping files (various sets of SNPs) have been uploaded to Dryad: https://doi.org/10.5061/dryad.j6q573ndp

https://briancoad.com

https://en.wikipedia.org/wiki/Tiber

## References

Aljanabi, S. M., & Martinez, I. (1997). Universal and rapid salt-extraction of high quality genomic DNA for PCR-based techniques. Nucleic acids research, 25(22), 4692–4693. doi.org/10.1093/nar/25.22.4692

Andrews, S., & Krueger, F. (2010). FastQC. A quality control tool for high throughput sequence data, 370.

Bandelt, H. J., Forster, P., & Röhl, A. (1999). Median-joining networks for inferring intraspecific phylogenies. Molecular Biology and Evolution, 16(1), 37–48.

Banister, K. E. (1984). A subterranean population of *Garra barreimiae* (Teleostei: Cyprinidae) from Oman, with comments on the concept of regressive evolution. Journal of Natural History, 18(6), 927–938.

Baumgarten, L., Machado-Schiaffino, G., Henning, F., & Meyer, A. (2015). What big lips are good for: on the adaptive function of repeatedly evolved hypertrophied lips of cichlid fishes. Biological Journal of the Linnean Society, 115(2), 448–455. doi.org/10.1111/bij.12502

Berg, L. S. (1914). Fishes. Vol. 3, Ostariophysi, Part. 2. St. Petersburg: Izd. Imper. Akad. Nauk (in Russian).

Benjamini, Y., & Hochberg, Y. (1995). Controlling the false discovery rate: a practical and powerful approach to multiple testing. Journal of the Royal statistical society: series B (Methodological), 57(1), 289–300.

Benvenuti, M., Carnicelli, S., Belluomini, G., Dainelli, N., Di Grazia, S., Ferrari, G. A., … Kebede, S. (2002). The Ziway–Shala lake basin (main Ethiopian rift, Ethiopia): a revision of basin evolution with special reference to the Late Quaternary. Journal of African Earth Sciences 35, 247–269.

Beshera, K. A., Harris, P. M., & Mayden, R. L. (2016). Novel evolutionary lineages in *Labeobarbus* (Cypriniformes; Cyprinidae) based on phylogenetic analyses of mtDNA sequences. Zootaxa, 4093(3), 363–381. doi.org/10.11646/zootaxa.4093.3.4

Braasch, I., & Postlethwait, J. H. (2012). Polyploidy in fish and the teleost genome duplication. In D. E. Soltis (Ed.), Polyploidy and genome evolution (pp. 341–383). Berlin, Heidelberg: Springer.

Brodersen, J., Post, D. M., & Seehausen, O. (2018). Upward adaptive radiation cascades: predator diversification induced by prey diversification. Trends in Ecology & Evolution, 33(1), 59–70. doi.org/10.1016/j.tree.2017.09.016

Burnashev, M. S. (1952). Snow trouts of the Zeravshan River. Proceedings of the Kishinev State University (Biology), 4, 111–125 (in Russian).

Burress, E. D., Piálek, L., Casciotta, J. R., Almirón, A., Tan, M., Armbruster, J. W., & Říčan, O. (2018). Island-and lake-like parallel adaptive radiations replicated in rivers. Proceedings of the Royal Society B: Biological Sciences, 285(1870), 20171762. doi.org/10.1098/rspb.2017.1762

Catchen, J., Hohenlohe, P. A., Bassham, S., Amores, A., & Cresko, W. A. (2013). Stacks: an analysis tool set for population genomics. Molecular Ecology, 22(11), 3124–3140. doi.org/10.1111/mec.12354

Chifman, J., & Kubatko, L. (2014). Quartet Inference from SNP Data Under the Coalescent Model. Bioinformatics, 30(23), 3317–3324, https://doi.org/10.1093/bioinformatics/btu530

Coad, B. W. (1996). Threatened fishes of the world: *Iranocypris typhlops* Bruun & Kaiser, 1944 (Cyprinidae). Environmental Biology of Fishes, 46(4), 374. https://doi.org/10.1007/BF00005015

Danecek, P., Auton, A., Abecasis, G., Albers, C. A., Banks, E., DePristo, M. A., … Durbin, R. (2011). The variant call format and VCFtools. Bioinformatics, 27(15), 2156–2158. doi: 10.1093/bioinformatics/btr330 doi.org/10.1093/bioinformatics/btr330 Data availability: https://doi.org/10.5061/dryad.j6q573ndp

de Graaf, M., Megens, H. J., Samallo, J., & Sibbing, F. (2007). Evolutionary origin of Lake Tana’s (Ethiopia) small *Barbus* species: indications of rapid ecological divergence and speciation. Animal Biology, 57(1), 39–48. doi.org/10.1163/157075607780002069

de Graaf, M., Megens, H. J., Samallo, J., & Sibbing, F. (2010). Preliminary insight into the age and origin of the *Labeobarbus* fish species flock from Lake Tana (Ethiopia) using the mtDNA cytochrome *b* gene. Molecular Phylogenetics and Evolution, 54(2), 336–343. doi.org/10.1016/j.ympev.2009.10.029

DeFaveri, J., & Merilä, J. (2013). Evidence for adaptive phenotypic differentiation in Baltic Sea sticklebacks. Journal of Evolutionary Biology, 26(8), 1700–1715. https://doi.org/10.1111/jeb.12168

Dejen, E., Rutjes, H. A., De Graaf, M., Nagelkerke, L. A., Osse, J. W., & Sibbing, F. A. (2002). The ‘small barbs’ *Barbus humilis* and *B. trispilopleura* of Lake Tana (Ethiopia): are they ecotypes of the same species?. Environmental Biology of Fishes, 65(4), 373–386. doi.org/10.1023/A:1021110721565

Dibaba, A., Soromessa, T., & Workineh, B., 2019. Carbon stock of the various carbon pools in Gerba-Dima moist Afromontane forest, South-western Ethiopia. Carbon Balance and Management, 14, 1. https://doi.org/10.1186/s13021-019-0116-x

Dimmick, W. W., Berendzen, P. B., & Golubtsov, A. S. (2001). Genetic comparison of three *Barbus* (Cyprinidae) morphotypes from the Genale River, Ethiopia. Copeia, 2001(4), 1123–1129. doi.org/10.1643/0045-8511(2001)001

Ewels, P., Magnusson, M., Lundin, S., & Käller, M. (2016). MultiQC: summarize analysis results for multiple tools and samples in a single report. Bioinformatics, 32(19), 3047–3048. doi.org/10.1093/bioinformatics/btw354

Feder, J. L., Egan, S. P., & Nosil, P. (2012). The genomics of speciation-with-gene-flow. Trends in Genetics, 28(7), 342–350. doi.org/10.1016/j.tig.2012.03.009

Ferguson, D. J., Barnie, T. D., Pyle, D. M., Oppenheimer, C., Yirgu, G., Lewi, E., … & Hamling, I. (2010). Recent rift-related volcanism in Afar, Ethiopia. Earth and Planetary Science Letters, 292(3-4), 409–418. doi.org/10.1016/j.epsl.2010.02.010

Feulner, P. G., Kirschbaum, F., & Tiedemann, R. (2008). Adaptive radiation in the Congo River: an ecological speciation scenario for African weakly electric fish (Teleostei; Mormyridae; *Campylomormyrus*). Journal of Physiology-Paris, 102(4-6), 340–346. doi.org/10.1016/j.jphysparis.2008.10.002

Franchini, P., Monné Parera, D., Kautt, A. F., & Meyer, A. (2017). quaddRAD: a new high-multiplexing and PCR duplicate removal ddRAD protocol produces novel evolutionary insights in a nonradiating cichlid lineage. Molecular Ecology, 26(10), 2783–2795. https://doi.org/10.1111/mec.14077

Fricke, R., Eschmeyer, W. N. & Van der Laan, R. (Eds.) 2021. ESCHMEYER’S CATALOG OF FISHES: GENERA, SPECIES, REFERENCES. (http://researcharchive.calacademy.org/research/ichthyology/catalog/fishcatmain.asp). Electronic version accessed 22 Feb 2021.

Fruciano, C., Franchini, P., Raffini, F., Fan, S., & Meyer, A. (2016). Are sympatrically speciating Midas cichlid fish special? Patterns of morphological and genetic variation in the closely related species *Archocentrus centrarchus*. Ecology and Evolution, 6(12), 4102–4114. https://doi.org/10.1002/ece3.2184

Fryer, G., & Iles, T. D. (1972). The Cichlid Fishes of the Great Lakes of Africa. Neptune City, NY: T.H.F. Publications Inc.

Geremew, A. (2007). Taxonomic Revision, Relative Abundance, and Aspects of the Biology of some Species of the Genus Garra, Hamilton 1922 (Pisces: Cyprinidae) in Lake Tana, Ethiopia (Unpublished doctoral dissertation). Addis Ababa University.

Getahun, A., & Stiassny, M. L. J., 1998. The freshwater biodiversity crisis: the case of the Ethiopian fish fauna. SINET: Ethiopian Journal of Science, 21, 207–230.

Givnish, T. J. (2015). Adaptive radiation versus ‘radiation’ and ‘explosive diversification’: why conceptual distinctions are fundamental to understanding evolution. New Phytologist, 207(2), 297–303. https://doi.org/10.1111/nph.13482

Glez-Peña, D., Gómez-Blanco, D., Reboiro-Jato, M., Fdez-Riverola, F., & Posada, D. (2010). ALTER: Program-oriented format conversion of DNA and protein alignments. Nucleic Acids Research, 38(Suppl 2), W14–W18. doi.org/10.1093/nar/gkq321.

Golubtsov, A. S. (2010). Fish ‘Species Flocks’ in Rivers and Lakes: Sympatric Divergence in Poor Fauna Fish Communities as Particular Modus of Evolution. In D. S. Pavlov, Y. Y. Dgebuadze & M. I. Shatunovsky (Eds.), Relevant Problems of Contemporary Ichthyology (To 100 Jubilee of G. V. Nikolsky) (pp. 96–123). Moscow: KMK Scientific Press.

Golubtsov, A. S., Cherenkov, S. E., & Tefera, F. (2012). High morphological diversity of the genus *Garra* in the Sore River (the White Nile Basin, Ethiopia): one more cyprinid species flock? Journal of Ichthyology, 52(11), 817–820. doi.org/10.1134/S0032945212110057

Golubtsov, A. S., & Darkov, A. A. 2008. A review of fish diversity in the main drainage systems of Ethiopia based on the data obtained by 2008. In D. S. Pavlov, Y. Y. Dgebuadze, A. A. Darkov, A. S. Golubtsov & M. V. Mina (Eds.), Ecological and faunistic studies in Ethiopia, Proceedings of jubilee meeting “Joint Ethio-Russian Biological Expedition (pp. 69–102). Moscow: KMK Scientific Press.

Golubtsov, A. S., Darkov, A. A., Dgebuadze, Y. Y., & Mina, M. V. (1995). An artificial key to fish species of the Gambela region (the White Nile basin in the limits of Ethiopia). Joint Ethio-Russian Biological Expedition. Addis Abeba.

Golubtsov, A. S., Dgebuadze, Y. Y., & Mina, M. V. (2002). Fishes of the Ethiopian Rift Valley. In C. Tudorancea & W. D. Taylor (Eds.), Ethiopian Rift Valley Lakes. Biology of Inland Waters Series (pp. 167–258). Leiden, The Netherlands: Backhuys Publishers.

Golubtsov, A. S., Korostelev, N. B., & Levin, B. A. (2021). Monsters with a shortened vertebral column: A population phenomenon in radiating fish *Labeobarbus* (Cyprinidae). Plos One, 16(1), e0239639. doi.org/10.1371/journal.pone.0239639

Hamidan, N., Jackson, M. C., & Britton, J. R. (2016). Diet and trophic niche of the endangered fish *Garra ghorensis* in three Jordanian populations. Ecology of Freshwater Fish, 25(3), 455–464. doi.org/10.1111/eff.12226

Hutchison, W., Fusillo, R., Pyle, D. M., Mather, T. A., Blundy, J. D., Biggs, J., … & Calvert, A. T. (2016). A pulse of mid-Pleistocene rift volcanism in Ethiopia at the dawn of modern humans. Nature Communications, 7(1), 1–12. doi.org/10.1038/ncomms13192

Hart, R. K., Calver, M. C., & Dickman, C. R. (2002). The index of relative importance: an alternative approach to reducing bias in descriptive studies of animal diets. Wildlife Research, 29(5), 415–421. doi.org/10.1071/WR02009

Henning, F., & Meyer, A. (2014). The evolutionary genomics of cichlid fishes: explosive speciation and adaptation in the postgenomic era. Annual Review of Genomics and Human Genetics, 15, 417–441. doi.org/10.1146/annurev-genom-090413-025412

Hoang, D. T., Chernomor, O., von Haeseler, A., Minh, B. Q., & Vinh, L. S. (2018). UFBoot2: Improving the Ultrafast Bootstrap Approximation. Molecular Biology and Evolution, 35(2), 518–522. doi.org/10.1093/molbev/msx281

Hubbs, C. L., & Lagler, K. F. (1958). Fishes of the Great Lakes region. Ann Arbor: Univ. Mich. Press.

Jacobs, A., Carruthers, M., Yurchenko, A., Gordeeva, N. V., Alekseyev, S. S., Hooker, O., … & Elmer, K. R. (2020). Parallelism in eco-morphology and gene expression despite variable evolutionary and genomic backgrounds in a Holarctic fish. PLoS Genetics, 16(4), e1008658. doi.org/10.1371/journal.pgen.1008658

Jombart, T. (2008). adegenet: a R package for the multivariate analysis of genetic markers. Bioinformatics, 24(11), 1403–1405. doi.org/10.1093/bioinformatics/btn129

Jombart, T., & Ahmed, I. (2011). adegenet 1.3-1: new tools for the analysis of genome-wide SNP data. Bioinformatics, 27(21), 3070–3071. doi.org/10.1093/bioinformatics/btr521

Joyce, D. A., Lunt, D. H., Bills, R., Turner, G. F., Katongo, C., Duftner, N., … & Seehausen, O. (2005). An extant cichlid fish radiation emerged in an extinct Pleistocene lake. Nature, 435(7038), 90–95. doi.org/10.1038/nature03489

Junker, J., Rick, J. A., McIntyre, P. B., Kimirei, I., Sweke, E. A., Mosille, J. B., … & Wagner, C. E. (2020) Structural genomic variation leads to genetic differentiation in Lake Tanganyika’s sardines. Molecular Ecology, 29: 3277–3298. https://doi.org/10.1111/mec.15559

Irisarri, I., Singh, P., Koblmüller, S., Torres-Dowdall, J., Henning, F., Franchini, P., … & Meyer, A. (2018). Phylogenomics uncovers early hybridization and adaptive loci shaping the radiation of Lake Tanganyika cichlid fishes. Nature communications, 9(1), 1–12. doi.org/10.1038/s41467-018-05479-9

Kalyaanamoorthy, S., Minh, B. Q., Wong, T. K., Von Haeseler, A., & Jermiin, L. S. (2017). ModelFinder: fast model selection for accurate phylogenetic estimates. Nature methods, 14(6), 587–589. doi.org/10.1038/nmeth.4285

Kautt, A. F., Machado-Schiaffino, G., & Meyer, A. (2016). Multispecies outcomes of sympatric speciation after admixture with the source population in two radiations of Nicaraguan crater lake cichlids. PLoS Genetics, 12(6), e1006157. doi.org/10.1371/journal.pgen.1006157

Kautt, A. F., Machado-Schiaffino, G., & Meyer, A. (2018). Lessons from a natural experiment: Allopatric morphological divergence and sympatric diversification in the Midas cichlid species complex are largely influenced by ecology in a deterministic way. Evolution Letters, 2(4), 323–340. doi.org/10.1002/evl3.64

Kautt, A. F., Kratochwil, C. F., Nater, A., Machado-Schiaffino, G., Olave, M., Henning, F., … & Meyer, A. (2020). Contrasting signatures of genomic divergence during sympatric speciation. Nature, 588(7836), 106–111. doi.org/10.1038/s41586-020-2845-0

Kebede, A., Diekkrüger, B., & Moges, S.A., 2014. Comparative study of a physically based distributed hydrological model versus a conceptual hydrological model for assessment of climate change response in the Upper Nile, Baro-Akobo basin: a case study of the Sore watershed, Ethiopia. International Journal of River Basin Management, 12(4), 299–318. http://dx.doi.org/10.1080/15715124.2014.917315

Kocher, T. D. (2004). Adaptive evolution and explosive speciation: the cichlid fish model. Nature Reviews Genetics, 5(4), 288–298. doi.org/10.1038/nrg1316

Kottelat, M. (2020). *Ceratogarra*, a genus name for *Garra cambodgiensis* and *G. fasciacauda* and comments on the oral and gular soft anatomy in labeonine fishes (Teleostei: Cyprinidae). The Raffles Bulletin of Zoology Supplement, 35, 156–178. DOI: 10.26107/RBZ-2020-0049

Kruckenhauser, L., Haring, E., Seemann, R., & Sattmann, H. (2011). Genetic differentiation between cave and surface-dwelling populations of *Garra barreimiae* (Cyprinidae) in Oman. BMC Evolutionary Biology, 11(1), 1–15. doi.org/10.1186/1471-2148-11-172

Krysanov, E. Y., & Golubtsov, A. S. (1993). Karyotypes of three *Garra* species from Ethiopia. Journal of fish biology, 42(3), 465–467.

Lanfear, R., Calcott, B., Ho, S. Y., & Guindon, S. (2012). PartitionFinder: combined selection of partitioning schemes and substitution models for phylogenetic analyses. Molecular Biology and Evolution, 29(6), 1695–1701. doi.org/10.1093/molbev/mss020

Langmead, B., & Salzberg, S. L. (2012). Fast gapped-read alignment with Bowtie 2. Nature Methods, 9(4), 357–359. doi: 10.1038/nmeth.1923 doi.org/10.1038/nmeth.1923

Lecaudey, L. A., Schliewen, U. K., Osinov, A. G., Taylor, E. B., Bernatchez, L., & Weiss, S. J. (2018). Inferring phylogenetic structure, hybridization and divergence times within Salmoninae (Teleostei: Salmonidae) using RAD-sequencing. Molecular Phylogenetics and Evolution, 124, 82–99. doi.org/10.1016/j.ympev.2018.02.022

Leigh, J. W., & Bryant, D. (2015). popart: full-feature software for haplotype network construction. Methods in Ecology and Evolution, 6(9), 1110–1116. doi.org/10.1111/2041-210X.12410

Levin, B.A., Casal-López, M., Simonov, E., Dgebuadze, Y.Y., Mugue, N.S., Tiunov, A.V., … Golubtsov, A.S. (2019). Adaptive radiation of barbs of the genus *Labeobarbus* (Cyprinidae) in an East African river. Freshwater Biology, 64, 1721–1736. https://doi.org/10.1111/fwb.13364

Levin, B.A., Simonov, E., Dgebuadze, Y.Y., Levina, M., & Golubtsov, A.S. (2020). In the rivers: Multiple adaptive radiations of cyprinid fishes (*Labeobarbus*) in Ethiopian Highlands. Scientific reports, 10(1), 7192. https://doi.org/10.1038/s41598-020-64350-4

Losos, J. B. (2010). Adaptive radiation, ecological opportunity, and evolutionary determinism: American Society of Naturalists EO Wilson Award address. The American Naturalist, 175(6), 623–639. DOI: 10.1086/652433

Machado-Schiaffino, G., Henning, F., & Meyer, A. (2014). Species-specific differences in adaptive phenotypic plasticity in an ecologically relevant trophic trait: hypertrophic lips in Midas cichlid fishes. Evolution, 68(7), 2086–2091. doi.org/10.1111/evo.12367

Machado-Schiaffino, G., Kautt, A. F., Torres-Dowdall, J., Baumgarten, L., Henning, F., & Meyer, A. (2017). Incipient speciation driven by hypertrophied lips in Midas cichlid fishes?. Molecular ecology, 26(8), 2348–2362. doi.org/10.1111/mec.14029

Malinsky, M., Svardal, H., Tyers, A. M., Miska, E. A., Genner, M. J., Turner, G. F., & Durbin, R. (2018) Whole-genome sequences of Malawi cichlids reveal multiple radiations interconnected by gene flow. Nature Ecology & Evolution, 457, 830. doi.org/10.1038/s41559-018-0717-x

Malinsky, M., Matschiner, M., & Svardal, H. (2021). Dsuite-fast D-statistics and related admixture evidence from VCF files. Molecular Ecology Resources, 21(2), 584–595. https://doi.org/10.1111/1755-0998.13265

Marques, D. A., Meier, J. I., & Seehausen, O. (2019). A combinatorial view on speciation and adaptive radiation. Trends in Ecology & Evolution, 34(6), 531–544. doi.org/10.1016/j.tree.2019.02.008

Martin, C. H., Cutler, J. S., Friel, J. P., Dening Touokong, C., Coop, G., & Wainwright, P. C. (2015). Complex histories of repeated gene flow in Cameroon crater lake cichlids cast doubt on one of the clearest examples of sympatric speciation. Evolution, 69(6), 1406–1422. doi.org/10.1111/evo.12674

Martin, M. (2011). Cutadapt removes adapter sequences from high-throughput sequencing reads. EMBnet.journal, 17(1), 10. doi: 10.14806/ej.17.1.200 doi.org/10.14806/ej.17.1.200

Matschiner, M., Hanel, R., & Salzburger, W. (2011). On the origin and trigger of the notothenioid adaptive radiation. PLoS One, 6(4), e18911. doi.org/10.1371/journal.pone.0018911

Matthes, H. (1963). A Comparative Study of the Feeding Mechanisms of Some African Cyprinidae (Pisces, Cypriniformes). Bijdragen tot de Dierkunde, 33(1), 3–24.

McKinnon, J. S., & Rundle, H. D. (2002). Speciation in nature: the threespine stickleback model systems. Trends in Ecology & Evolution, 17(10), 480–488. doi.org/10.1016/S0169-5347(02)02579-X

Mège, D., Purcell, P., Pochat, S., & Guidat, T. (2015). The landscape and landforms of the Ogaden, Southeast Ethiopia. In P. Billi (Ed.), Landscapes and landforms of Ethiopia (pp. 323–348). Dordrecht, The Netherlands: Springer.

Meier, J. I., Marques, D. A., Mwaiko, S., Wagner, C. E., Excoffier, L., & Seehausen, O. (2017). Ancient hybridization fuels rapid cichlid fish adaptive radiations. Nature communications, 8(1), 1–11. doi.org/10.1038/ncomms14363

Menon, A. G. K. (1964). Monograph of the cyprinid fishes of the genus Garra Hamilton (Vol. 14). Government of India.

Melaku, S., Abebe Getahun, A., & Wakjira, M. (2017). Population aspects of fishes in Geba and Sor rivers, White Nile System in Ethiopia, East Africa. International Journal of Biodiversity, 2017, 1252604. https://doi.org/10.1155/2017/1252604

Melnik, N. O., Markevich, G. N., Taylor, E. B., Loktyushkin, A. V., & Esin, E. V. (2020). Evidence for divergence between sympatric stone charr and Dolly Varden along unique environmental gradients in Kamchatka. Journal of Zoological Systematics and Evolutionary Research, 58(4), 1135–1150. doi.org/10.1111/jzs.12367

Meyer, A., Kocher, T. D., Basasibwaki, P., & Wilson, A. C. (1990). Monophyletic origin of Lake Victoria cichlid fishes suggested by mitochondrial DNA sequences. Nature, 347(6293), 550–553.

Mina, M. V., Levin, B. A., & Mironovsky, A. N. (2005). On the possibility of using character estimates obtained by different operators in morphometric studies of fish. Journal of Ichthyology, 45(4), 284–294.

Mina, M. V., Mironovsky, A. N., & Dgebuadze, Y. (1996). Lake Tana large barbs: phenetics, growth and diversification. Journal of Fish Biology, 48(3), 383–404.

Mina, M. V., Mironovsky, A. N., Golubtsov, A. S., & Dgebuadze, Y. Y. (1998). II. Morphological diversity of “large barbs”; from Lake Tana and neighbouring areas: Homoplasies or synapomorphies?. Italian Journal of Zoology, 65(S1), 9–14.

Minh, B. Q., Schmidt, H. A., Chernomor, O., Schrempf, D., Woodhams, M. D., Von Haeseler, A., & Lanfear, R. (2020). IQ-TREE 2: New models and efficient methods for phylogenetic inference in the genomic era. Molecular Biology and Evolution, 37(5), 1530–1534. https://doi.org/10.1093/molbev/msaa015

Mironovsky, A. N., Mina, M. V., & Dgebuadze, Y. Y. (2019). Large African Barbs with Hypertrophied Lips and their Relationship with Generalized Forms of Species of the Genus *Barbus* (*Labeobarbus* auctorum). Journal of Ichthyology, 59(3), 327–335. doi.org/10.1134/S0032945219030111

Moritz, T., El Dayem, Z.N., Abdallah, M.A., & Neumann, D. (2019). New and rare records of fishes from the White Nile in the Republic of the Sudan. Cybium, 43, 137–151. https://doi.org/10.26028/cybium/2019-423-011

Mousavi-Sabet, H., & Eagderi, S. (2016). *Garra lorestanensis*, a new cave fish from the Tigris River drainage with remarks on the subterranean fishes in Iran (Teleostei: Cyprinidae). FishTaxa, 1(1), 45–54. http://dx.doi.org/10.7508/jft.2016.01.006

Muschick, M., Nosil, P., Roesti, M., Dittmann, M. T., Harmon, L., & Salzburger, W. (2014). Testing the stages model in the adaptive radiation of cichlid fishes in East African Lake Tanganyika. Proceedings of the Royal Society B: Biological Sciences, 281(1795), 20140605. doi.org/10.1098/rspb.2014.0605

Nagelkerke, L. (1997). The barbs of Lake Tana, Ethiopia: morphological diversity and its implications for taxonomy, trophic resource partitioning, and fisheries (Unpublished doctoral dissertation). Agricultural University of Wageningen.

Nagelkerke, L. A., Sibbing, F. A., van den Boogaart, J. G., Lammens, E. H., & Osse, J. W. (1994). The barbs (*Barbus* spp.) of Lake Tana: a forgotten species flock?. Environmental Biology of Fishes, 39(1), 1–22.

Nagelkerke, L. A. J., & Sibbing, F. A. (1997). A revision of the large barbs (Barbus spp., Cyprinidae, Teleostei) of Lake Tana, Ethiopia, with a description of seven new species. In: The barbs of Lake Tana, Ethiopia: morphological diversity and its implications for taxonomy, trophic resource partitioning, and fisheries (pp. 105–170). (Unpublished doctoral dissertation). Agricultural University of Wageningen.

Nagelkerke, L. A. J., Leon-Kloosterziel, K. M., Megens, H. J., De Graaf, M., Diekmann, O. E., & Sibbing, F. A. (2015). Shallow genetic divergence and species delineations in the endemic *Labeobarbus* species flock of Lake Tana, Ethiopia. Journal of Fish Biology, 87(5), 1191–1208. doi.org/10.1111/jfb.12779

Neumann, D., Obermaier, H., & Moritz, T. 2016. Annotated checklist for fishes of the Main Nile Basin in the Sudan and Egypt based on recent specimen records (2006-2015). Cybium, 40: 287–317. doi.org/10.26028/cybium/2016-404-004

Nguyen, L. T., Schmidt, H. A., Von Haeseler, A., & Minh, B. Q. (2015). IQ-TREE: a fast and effective stochastic algorithm for estimating maximum-likelihood phylogenies. Molecular Biology and Evolution, 32(1), 268–274. doi.org/10.1093/molbev/msu300

Østbye, K., Amundsen, P. A., Bernatchez, L., Klemetsen, A., Knudsen, R., Kristoffersen, R., … & Hindar, K. (2006). Parallel evolution of ecomorphological traits in the European whitefish *Coregonus lavaretus* (L.) species complex during postglacial times. Molecular Ecology, 15(13), 3983–4001. doi.org/10.1111/j.1365-294X.2006.03062.x

Palumbi, S. R. (1996). Nucleic acids II: The polymerase chain reaction. In D. M. Hillis, C. Moritz & B. K. Mable (Eds.), Molecular systematics (pp. 205–247). Sunderland, MA: Sinauer Associates.

Paris, J. R., Stevens, J. R., & Catchen, J. M. (2017). Lost in parameter space: a road map for stacks. Methods in Ecology and Evolution, 8(10), 1360–1373. doi.org/10.1111/2041-210X.12775

Patterson, N., Moorjani, P., Luo, Y., Mallick, S., Rohland, N., Zhan, Y., … & Reich, D. (2012). Ancient admixture in human history. Genetics, 192(3), 1065–1093. doi.org/10.1534/genetics.112.145037

Peichel, C. L., Nereng, K. S., Ohgi, K. A., Cole, B. L., Colosimo, P. F., Buerkle, C. A., … & Kingsley, D. M. (2001). The genetic architecture of divergence between threespine stickleback species. Nature, 414(6866), 901–905. doi.org/10.1038/414901a

Perdices, A., & Doadrio, I. (2001). The molecular systematics and biogeography of the European cobitids based on mitochondrial DNA sequences. Molecular Phylogenetics and Evolution, 19(3), 468–478. doi.org/10.1006/mpev.2000.0900

Piálek, L., Říčan, O., Casciotta, J., Almirón, A., & Zrzavý, J. (2012). Multilocus phylogeny of *Crenicichla* (Teleostei: Cichlidae), with biogeography of the *C. lacustris* group: species flocks as a model for sympatric speciation in rivers. Molecular Phylogenetics and Evolution, 62(1), 46–61. doi.org/10.1016/j.ympev.2011.09.006

Præbel, K., Knudsen, R., Siwertsson, A., Karhunen, M., Kahilainen, K. K., Ovaskainen, O., … & Amundsen, P. A. (2013). Ecological speciation in postglacial European whitefish: rapid adaptive radiations into the littoral, pelagic, and profundal lake habitats. Ecology and Evolution, 3(15), 4970–4986. doi.org/10.1002/ece3.867

Prave, A. R., Bates, C. R., Donaldson, C. H., Toland, H., Condon, D. J., Mark, D., & Raub, T. D. (2016). Geology and geochronology of the Tana Basin, Ethiopia: LIP volcanism, super eruptions and Eocene–Oligocene environmental change. Earth and Planetary Science Letters, 443, 1–8. doi.org/10.1016/j.epsl.2016.03.009

Prokofiev, A. M. & Golubtsov A. S. (2013). Revision of the loach genus *Afronemacheilus* (Teleostei: Balitoridae: Nemacheilinae) with description of a new species from the Omo-Turkana basin, Ethiopia. Ichthyological Exploration of Freshwaters, 24, 1–14.

Puebla, O. (2009). Ecological speciation in marine v. freshwater fishes. Journal of Fish Biology, 75(5), 960–996. doi.org/10.1111/j.1095-8649.2009.02358.x

Rambaut, A. (2014). FigTree 1.4.2 software. Institute of Evolutionary Biology, Univ. Edinburgh.

Rambaut, A., Ho, S. Y., Drummond, A. J., & Shapiro, B. (2009). Accommodating the effect of ancient DNA damage on inferences of demographic histories. Molecular Biology and Evolution, 26(2), 245–248. doi.org/10.1093/molbev/msn256

Rambaut, A., Suchard, M. A., Xie, D. & Drummond, A. J. (2014). Tracer v1.6. Retrieved from http://beast.bio.ed.ac.uk/Tracer

Reich, D., Thangaraj, K., Patterson, N., Price, A. L., & Singh, L. (2009). Reconstructing Indian population history. Nature, 461(7263), 489–494. doi.org/10.1038/nature08365

Ribbink, A. J., Marsh, A. C., Marsh, B. A., & Sharp, B. J. (1983). The zoogeography, ecology and taxonomy of the genus *Labeotropheus* Ahl, 1927, of Lake Malawi (Pisces: Cichlidae). Zoological Journal of the Linnean Society, 79(3), 223–243.

Richards, E. J., Servedio, M. R., & Martin, C. H. (2019). Searching for sympatric speciation in the genomic era. BioEssays, 41(7), 1900047. doi.org/10.1002/bies.201900047

Roberts, T. R. (1998). Review of the tropical Asian cyprinid fish genus *Poropuntius*, with descriptions of new species and trophic morphs. Natural History Bulletin of the Siam Society, 46(1), 105–135.

Roberts, T. R., & Khaironizam, M. Z. (2008). Trophic polymorphism in the Malaysian fish *Neolissochilus soroides* and other old world barbs (Teleostei, Cyprinidae). Natural History Bulletin of the Siam Society, 56, 25–53.

Rougeux, C., Bernatchez, L., & Gagnaire, P. A. (2017). Modeling the multiple facets of speciation-with-gene-flow toward inferring the divergence history of lake whitefish species pairs (*Coregonus clupeaformis*). Genome Biology and Evolution, 9(8), 2057–2074. doi.org/10.1093/gbe/evx150

Ronquist, F., Teslenko, M., Van Der Mark, P., Ayres, D. L., Darling, A., Höhna, S., … & Huelsenbeck, J. P. (2012). MrBayes 3.2: efficient Bayesian phylogenetic inference and model choice across a large model space. Systematic Biology, 61(3), 539–542. doi.org/10.1093/sysbio/sys029

Salzburger, W., Mack, T., Verheyen, E., & Meyer, A. (2005). Out of Tanganyika: genesis, explosive speciation, key-innovations and phylogeography of the haplochromine cichlid fishes. BMC Evolutionary Biology, 5(1), 1–15. doi.org/10.1186/1471-2148-5-17

Schluter, D. (2000). The ecology of adaptive radiation. New York: Oxford University Press.

Schwarzer, J., Misof, B., Ifuta, S. N., & Schliewen, U. K. (2011). Time and origin of cichlid colonization of the lower Congo rapids. Plos One, 6(7), e22380. doi.org/10.1371/journal.pone.0022380

Seehausen, O. (2000). Explosive speciation rates and unusual species richness in haplochromine cichlid fishes: effects of sexual selection. Advances in Ecological Research, 31, 237–274. doi.org/10.1016/S0065-2504(00)31015-7

Seehausen, O. (2006). African cichlid fish: a model system in adaptive radiation research. Proceedings of the Royal Society B: Biological Sciences, 273(1597), 1987–1998. doi.org/10.1098/rspb.2006.3539

Seehausen, O., & Wagner, C. E. (2014). Speciation in freshwater fishes. Annual Review of Ecology, Evolution, and Systematics, 45, 621–651. doi.org/10.1146/annurev-ecolsys-120213-091818

Segherloo, I. H., Normandeau, E., Benestan, L., Rougeux, C., Coté, G., Moore, J. S., … & Bernatchez, L. (2018). Genetic and morphological support for possible sympatric origin of fish from subterranean habitats. Scientific Reports, 8(1), 1–13. doi.org/10.1038/s41598-018-20666-w

Selz, O. M., & Seehausen, O. (2019). Interspecific hybridization can generate functional novelty in cichlid fish. Proceedings of the Royal Society B, 286(1913), 20191621. doi.org/10.1098/rspb.2019.1621

Sibbing, F. A., Nagelkerke, L. A., Stet, R. J., & Osse, J. W. (1998). Speciation of endemic Lake Tana barbs (Cyprinidae, Ethiopia) driven by trophic resource partitioning; a molecular and ecomorphological approach. Aquatic Ecology, 32(3), 217–227.

Skúlason, S. (1999). Sympatric morphs, populations and speciation in freshwater fish with emphasis on arctic charr. In A. Magurran & R. M. May (Eds.), Evolution of biological diversity (pp. 71–92). New York: Oxford University Press.

Smadja, C. M., & Butlin, R. K. (2011). A framework for comparing processes of speciation in the presence of gene flow. Molecular Ecology, 20(24), 5123–5140. doi.org/10.1111/j.1365-294X.2011.05350.x

Stiassny, M.L.J. & Abebe Getahun. 2007. An overview of labeonin relationships and the phylogenetic placement of the Afro-Asian genus *Garra* Hamilton, 1822 (Teleostei: Cyprinidae), with the description of five new species of *Garra* from Ethiopia, and a key to all African species. Zoological Journal of the Linnean Society, 150, 41–83. doi.org/10.1111/j.1096-3642.2007.00281.x

Stewart, K. M., & Murray, A. M. (2017). Biogeographic implications of fossil fishes from the Awash River, Ethiopia. Journal of Vertebrate Paleontology, 37(1), e1269115. doi.org/10.1080/02724634.2017.1269115

Sturmbauer, C. (1998). Explosive speciation in cichlid fishes of the African Great Lakes: a dynamic model of adaptive radiation. Journal of Fish Biology, 53, 18–36. doi.org/10.1111/j.1095-8649.1998.tb01015.x

Swofford, D. L. 2003. PAUP*. Phylogenetic Analysis Using Parsimony (and Other Methods). Version 4. Sinauer Associates, Sunderland, Massachusetts.

Taylor, E. B. (1999). Species pairs of north temperate freshwater fishes: evolution, taxonomy, and conservation. Reviews in Fish Biology and Fisheries, 9(4), 299–324. doi.org/10.1023/A:1008955229420

Tamura, K., Stecher, G., Peterson, D., Filipski, A., & Kumar, S. (2013). MEGA6: molecular evolutionary genetics analysis version 6.0. Molecular Biology and Evolution, 30(12), 2725–2729. doi.org/10.1093/molbev/mst197

Tang, Q., Getahun, A., & Liu, H. (2009). Multiple in-to-Africa dispersals of labeonin fishes (Teleostei: Cyprinidae) revealed by molecular phylogenetic analysis. Hydrobiologia, 632(1), 261–271. doi.org/10.1007/s10750-009-9848-z

Terashima, A. (1984). Three new species of the cyprinid genus *Schizothorax* from Lake Rara, northwestern Nepal. Japanese Journal of Ichthyology, 31(2), 122–135.

Terekhanova, N. V., Logacheva, M. D., Penin, A. A., Neretina, T. V., Barmintseva, A. E., Bazykin, G. A., … & Mugue, N. S. (2014). Fast evolution from precast bricks: genomics of young freshwater populations of threespine stickleback *Gasterosteus aculeatus*. PLoS Genetics, 10(10), e1004696. doi.org/10.1371/journal.pgen.1004696

Verheyen, E., Salzburger, W., Snoeks, J., & Meyer, A. (2003). Origin of the superflock of cichlid fishes from Lake Victoria, East Africa. Science, 300(5617), 325–329. DOI:10.1126/science.1080699

Verity, R., & Nichols, R. A. (2016). Estimating the Number of Subpopulations (K) in Structured Populations. Genetics, 203(4), 1827–1839. doi:10.1534/genetics.115.180992

Vreven, E. J., Musschoot, T., Snoeks, J., & Schliewen, U. K. (2016). The African hexaploid Torini (Cypriniformes: Cyprinidae): review of a tumultuous history. Zoological Journal of the Linnean Society, 177(2), 231–305. doi.org/10.1111/zoj.12366

Wagner, C. E., Harmon, L. J., & Seehausen, O. (2012). Ecological opportunity and sexual selection together predict adaptive radiation. Nature, 487(7407), 366–369. doi.org/10.1038/nature11144

Wagner, C. E., McIntyre, P. B., Buels, K. S., Gilbert, D. M., & Michel, E. (2009). Diet predicts intestine length in Lake Tanganyika’s cichlid fishes. Functional Ecology, 23(6), 1122–1131. doi.org/10.1111/j.1365-2435.2009.01589.x

Whiteley, A. R. (2007). Trophic polymorphism in a riverine fish: morphological, dietary, and genetic analysis of mountain whitefish. Biological Journal of the Linnean Society, 92(2), 253–267. doi.org/10.1111/j.1095-8312.2007.00845.x

Yang, L., Arunachalam, M., Sado, T., Levin, B. A., Golubtsov, A. S., Freyhof, J., … & He, S. (2012). Molecular phylogeny of the cyprinid tribe Labeonini (Teleostei: Cypriniformes). Molecular Phylogenetics and Evolution, 65(2), 362–379. doi.org/10.1016/j.ympev.2012.06.007

Zandonà, E., Auer, S. K., Kilham, S. S., & Reznick, D. N. (2015). Contrasting population and diet influences on gut length of an omnivorous tropical fish, the Trinidadian guppy (*Poecilia reticulata*). PLoS One, 10(9), e0136079. doi.org/10.1371/journal.pone.0136079

Zheng, X. M., & Ge, S. (2010). Ecological divergence in the presence of gene flow in two closely related Oryza species (*Oryza rufipogon* and *O. nivara*). Molecular Ecology, 19(12), 2439–2454. doi.org/10.1111/j.1365-294X.2010.04674.x

